# Structural dynamics underlying agonist activation of a GLP-1R-Gs precoupled complex

**DOI:** 10.64898/2026.08.03.742488

**Authors:** Jakub Sýs, Joseph D Ho, Aaron D Showalter, An-Ping Yu, David B Wainscott, Veronica Laos, Howard Broughton, Kyle W Sloop, Alfonso Espada, Eamonn Reading

## Abstract

G-protein-coupled receptors (GPCRs) act as allosteric transmembrane signalling machines, generating distinct cellular responses depending on the conformational states induced by ligand binding. The glucagon-like peptide-1 receptor (GLP-1R), a class B GPCR central to insulin secretion and body-weight regulation, is a key therapeutic target for obesity-associated metabolic disease. Here, we used hydrogen–deuterium exchange mass spectrometry to characterize ligand-evoked structural dynamics within a pre-coupled GLP-1R–Gs protein complex. Non-peptide agonists Chu-128 and danuglipron elicited overlapping dynamic perturbation profiles, with distinct drug-specific effects within the transmembrane bundle. In contrast, the natural GLP-1 hormone produced a weaker stabilizing effect on receptor backbone dynamics, while its inactive metabolite exerted opposing localised destabilization. Notably, both peptides uniquely modulated the highly flexible G-protein switch III loop, a key mediator of downstream signalling. These findings pinpoint areas where structural dynamics shape agonist efficacy and facilitate functional dynamics-integrated drug discovery of non-peptide agonists.

**Significance:** The development of non-peptide agonists of the glucagon-like peptide-1 receptor (GLP-1R) represents a major advance in metabolic therapeutics, addressing key limitations of current peptide-based incretin therapies while enabling improved control over receptor signalling and pharmacokinetic properties. The recent FDA approval of a first-in-class small-molecule oral GLP-1R agonist, LY3502970, highlights the translational potential of this approach. However, the molecular basis by which distinct ligands modulate GLP-1R conformational dynamics and signalling remains poorly understood. Here we report how non-peptide agonists (Chu-128 and danuglipron) and endogenous GLP-1 peptide and its inactive metabolite shape the structural dynamics of a pre-coupled GLP-1R–Gs complex, revealing patterns linked to receptor activation. Our findings provide insights that guide the rational design of next-generation GLP-1R therapeutics.

## Introduction

G-protein coupled receptors (GPCRs) are transmembrane proteins which regulate a broad variety of functions in our body from vital functions like homeostasis through to vision, learning and memory (1). There are six recognised GPCR classes (2, 3) from which the B class is of major interest in treatment of endocrine diseases and metabolic conditions. During the past two decades, increasing attention has been directed toward members of class B for treatment of type 2 diabetes mellitus (T2DM) and obesity therapy (4, 5), including the glucagon-like peptide 1 receptor (GLP-1R), which is involved in glucose homeostasis (6). GLP-1 is an incretin peptide whose binding to GLP-1R induces activation of heterotrimeric stimulatory G-protein (G_s_) coupled to the receptor, which in turn potentiates protein kinase A responsible for producing cAMP (7). The cAMP-sensitive ion channels are then activated (8) which leads to glucose-dependent insulin secretion from pancreatic β-cells (9–11). GLP-1R is also active within pancreatic *α*- and δ-cells where GLP-1 positively lowers blood glucose via inhibition of glucagon secretion. Furthermore, GLP-1R agonism inhibits gastric emptying and activates anorexigenic pathways in the central nervous system that promote satiety (12–15). These receptor functions underpin why GLP-1R has become a principal therapeutic target in the treatment of T2DM and obesity-management.

Endogenous GLP-1 (GLP-1[7-36]) is rapidly cleaved by dipeptidyl peptidase-4 (DPP-4), within 2 minutes, to form the GLP-1[9-36] metabolite, leading to its degradation in human serum. Therefore, due to this rapid processing, therapeutic utilisation of GLP-1 is nearly impractical (16, 17). Given that, peptide engineering efforts were undertaken to develop more stable GLP- 1 mimetics as therapeutics, ultimately supporting obesity and weight-loss management. As early as 2026, only peptide GLP-1 mimetics like semaglutide (18, 19), liraglutide (20, 21), and tirzepatide (22, 23) have been approved. Although several non-peptide GLP-1 mimicking agonists entered clinical trials, the development of agents like PF-06882961 (danuglipron; 24) or taspoglutide has been discontinued and their clinical trial has been halted due to safety concerns (25). To date, the orally administered non-peptide GLP-1 mimetic LY3502970 (Orforglipron; 26) is the only such molecule approved by a regulatory authority, as it recently received FDA approval for use as an effective anti-obesity medication (27). LY3502970 displays some similar features to the Chu-128 analogue, a heavily Gs protein-biased GLP-1R non-peptide agonist, which previously showed it has different pharmacological profiles to danuglipron (28). Moreover, Orforglipron was shown to share Chu-128-induced cAMP production with even significantly reduced β-arrestin recruitment (29). However, reasons why partial agonist Orforglipron is capable to outperform full-agonist danuglipron in cAMP production and safety feature delivering stronger weight reduction are still unclear.

Structurally GLP-1R possesses the archetypal GPCR seven transmembrane (7TM) helical bundle but is distinguished by its large, specialised N-terminal extracellular domain (ECD) and lack of “DRY ionic-lock” motif found in TM3 of class A receptors (replacing it instead with conserved HETx and Pro-X-X-Gly (PxxG) motifs). Whilst also exhibiting a sharp kink and disruption of the helical fold in TM6. GPCRs exhibit intrinsic allosteric structural plasticity, enabling them to adopt multiple conformations and engage diverse downstream effector proteins to modulate cell signalling (30–32). It is becoming increasingly appreciated that comprehensive insight into their structural and functional dynamics (33, 34) is important to elucidate mechanisms driving GPCR activity, and shed light into GPCRs’ biased signalling, a phenomenon which is poorly understood. With recent evidence providing insight into canonical and non-canonical intermediate states of G-protein engagement within GPCRs, which are proposed to represent intermediate forms along the activation pathway of G proteins (35, 36).

Here, we used hydrogen–deuterium exchange mass spectrometry (HDX-MS) to uncover the structural dynamics that coordinate GLP-1R G_s_-protein (stimulatory G-protein, consisting of the Gα and the tightly associated Gβγ subunits) signalling by both peptide, GLP-1[7-36] and GLP-1[9-36], and non-peptide agonists, Chu-128 and danuglipron (**Fig. 1A-B**; 37). In an HDX-MS experiment, a protein system is exposed to D₂O solvent, allowing backbone amide hydrogens to exchange with deuterium at rates that reflect local and global structural dynamics. The degree of exchange is subsequently extracted following D-label quenching, protein digestion into peptides and LC-MS analysis and data processing (38, 39). HDX-MS reports on the solution-phase equilibrium of opening rates of backbone amide protons (EX2 kinetics) but is also capable of detecting the presence of kinetically distinct populations through identification of spectral multimodality (so-called EX1 or EXX kinetics) (40, 41). Bimodality has been discovered previously in GPCRs and are of particular interest for understanding activation mechanisms, being observed at the C-terminus of the TM6/ECL3 region of the β2- adrenergic receptor and in several regions of the glucagon receptor including the extracellular domain, TM1, TM2 and TM6 (42, 43).

**Figure 1.**
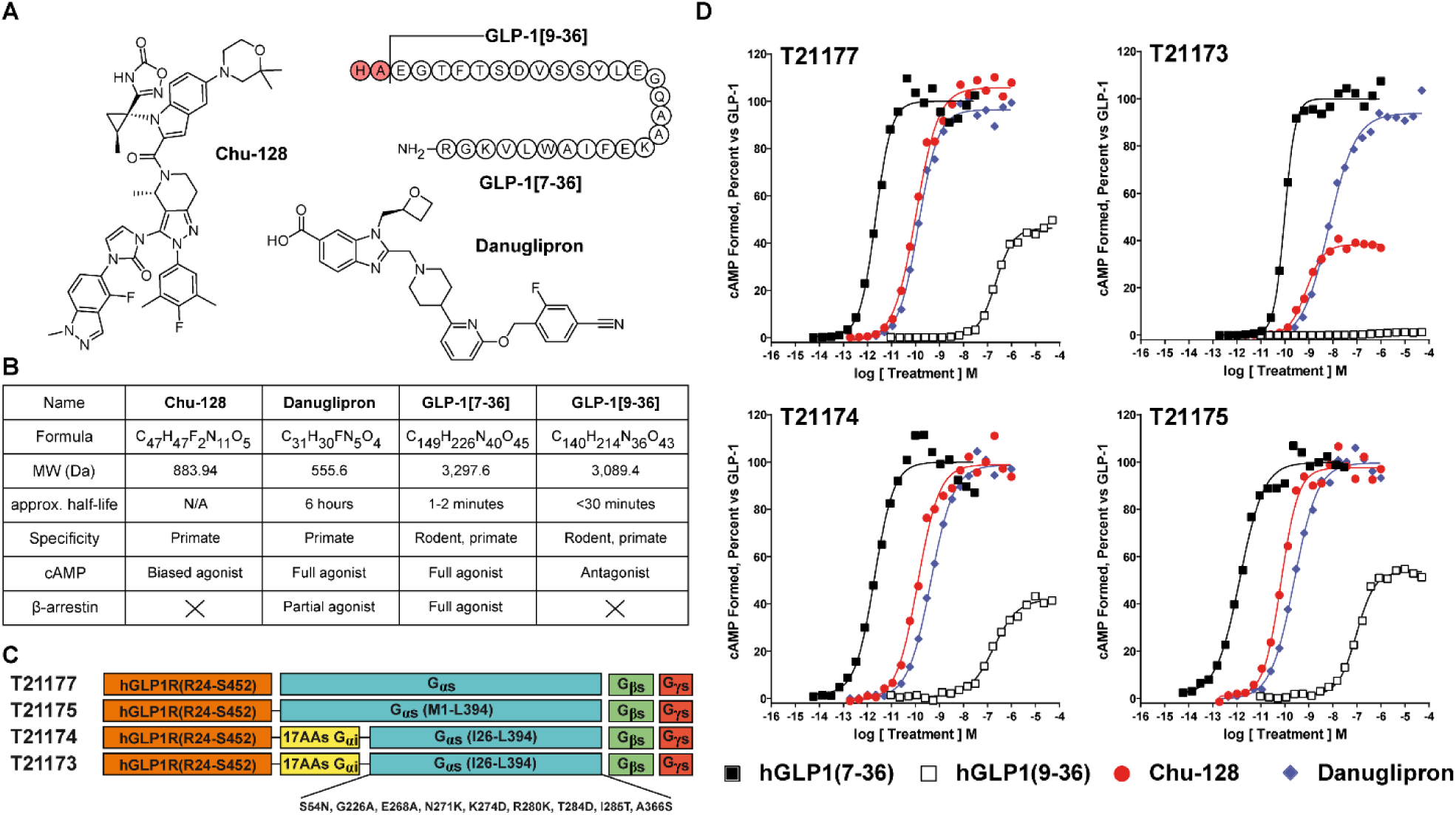
Functional Characterisation of Pre-Coupled GLP-1R–Gs via cAMP Production. (**A**) Chemical structures of ligands used in study and (**B**) their characteristics: Chu-128 and danuglipron non-peptide agonists (NPAs), and GLP-1[7-36] and GLP-1[9-36] peptides. **(C)** Linear representation of primary structure of human GLP-1R (hGLP-1R) constructs tested within cAMP functional assay. (**D**) Functional assays measuring cAMP accumulation in response to ligand activation. Constructs tested: T21177 = hGLP1R(R24-S452), T21173 (construct used for study, with mutations favouring nucleotide-free state) = hGLP1R(R24- S452) - (17 AAs from G-alpha-i) - G-alpha-s (I26-L394) (S54N, G226A, E268A, N271K, K274D, R280K, T284D, I285T, A366S), T21174 = hGLP1R(R24-S452) - (17 AAs from G- alpha-i) - G-alpha-s (I26-L394), T21175 = hGLP1R(R24-S452) - G-alpha-s(M1-L394)). Mutation favouring nucleotide-free state of hGLP1R(R24-S452).

In this work, we reveal that non-peptide agonists (danuglipron, Chu-128) broadly stabilise GLP- 1R receptor backbone dynamics and conformational switching with ligand-specific regional differences, while GLP-1 peptides likely act mainly via side-chain interaction modifications. Despite these differences, all agonists stabilise the receptor–G_s_ protein complex. In contrast, GLP-1[9–36] was found to promote a more dynamic receptor conformation than GLP-1[7–36] or NPAs. Collectively, these results provide mechanistic insights into GLP-1R functional dynamics that could guide the development of improved GLP-1R therapeutics.

## Results

### Capturing structural dynamics of pre-coupled GLP-1R-Gs

For HDX-MS to be most informative on GLP-1R function, acquisition of a ligand-free state is optimal to derive conclusions on the dynamical alterations between apo and ligand-bound states. Moreover, capture of a GLP-1R G_s_-protein complex coupling provides mechanistic insight into GLP-1R-Gs signalling. Initially, we attempted to purify a ligand-free noncovalent GLP-1R:G_s_ complex without any stabilising modifications and in absence of agonist, following previously established protocols successful for purification of the ligand-bound form (29, 44). In the absence of ligand, purification via Lauryl Maltose Neopentyl Glycol/Cholesteryl hemisuccinate (LMNG/CHS) solubilisation and FLAG-tag immobilisation (positioned at GLP- 1R N-terminus) resulted in low purity and GLP-1R partially truncated (*see* **Supplementary Discussion** and **Fig. S1A-E**).

To achieve a stable ligand-free GLP-1R coupled to Gs-protein we developed a construct where GLP-1R was recombinantly linked to the N-terminus of the Gα protein (herein denoted as GLP- 1R-Gs), with mutations made in Gα to favour a nucleotide-free state commonly used to provide a stable, high-affinity, active GPCR and G-protein complex state (**Fig. 1C** and **Fig. S2**; 45). Recombinant-linking of Gα to GLP-1R did not significantly alter cAMP accumulation profiles for the ligands investigated in this study: Chu-128 and danuglipron non-peptide agonists (NPAs), and GLP-1[7-36], the natural agonist of GLP-1R, and its DPP-4 degraded metabolite GLP-1[9-36] (**Fig. 1D**). Whereas the mutations to favour a nucleotide-free state reduced cAMP accumulation extent due to anticipated reduced nucleotide turnover. Radioligand competition binding assays further validated the affinities of the NPAs and peptides GLP-1[7-36] and GLP- 1[9-36] for the purified detergent-solubilized and native membrane-embedded protein complexes (**Table S1**). Overall, this provided a tractable pre-coupled nucleotide-free GLP-1R- Gs complex for structural biology investigation, but it is important to note it may contain structural changes in G protein binding that are not observed in the transient nucleotide-free state *in vivo*.

Through optimisation of HDX-MS experimental quench, digestion and LC-MS conditions, we were able to identify 211 peptides covering 78.6% of the entire sequence of the GLP-1R-Gs construct (protein-specific sequence coverage: GLP-1R, 67.2%; G_α_, 95.6%; G_β_, 93.9%) with an average redundancy of 3.79 (**Fig. 2**). We could not identify any peptides spanning extracellular loop 2 (ECL2), intracellular loops 1 and 2 (ICL1 and ICL2), and transmembrane helix 3 (TM3) in GLP-1R, as well as for the small 7-8 kDa Gγ subunit. This is likely due to the known difficulties in full digestion of hydrophobic TM domains and digestion pattern driven by specificity of co-immobilised pepsin and nepenthesin-II proteases (46). Achieving sufficient sequence coverage, we then performed HDX labelling at 10, 50, 1,250 (20 min 50 s), and 6,250 seconds (1 h 44 min 10 s), enabling us to characterise the dynamic behaviour of GLP-1R-Gs in its ligand-free state. To visualize the HDX data, we used AlphaFold 3 to generate a receptor– Gs complex model, enabling mapping of regions unresolved in available cryo-EM structures (**Fig. S3-4**). Where possible, data were mapped onto experimentally solved GLP-1R structures bound to Chu-128 (PDB:6X19).

**Figure 2.**
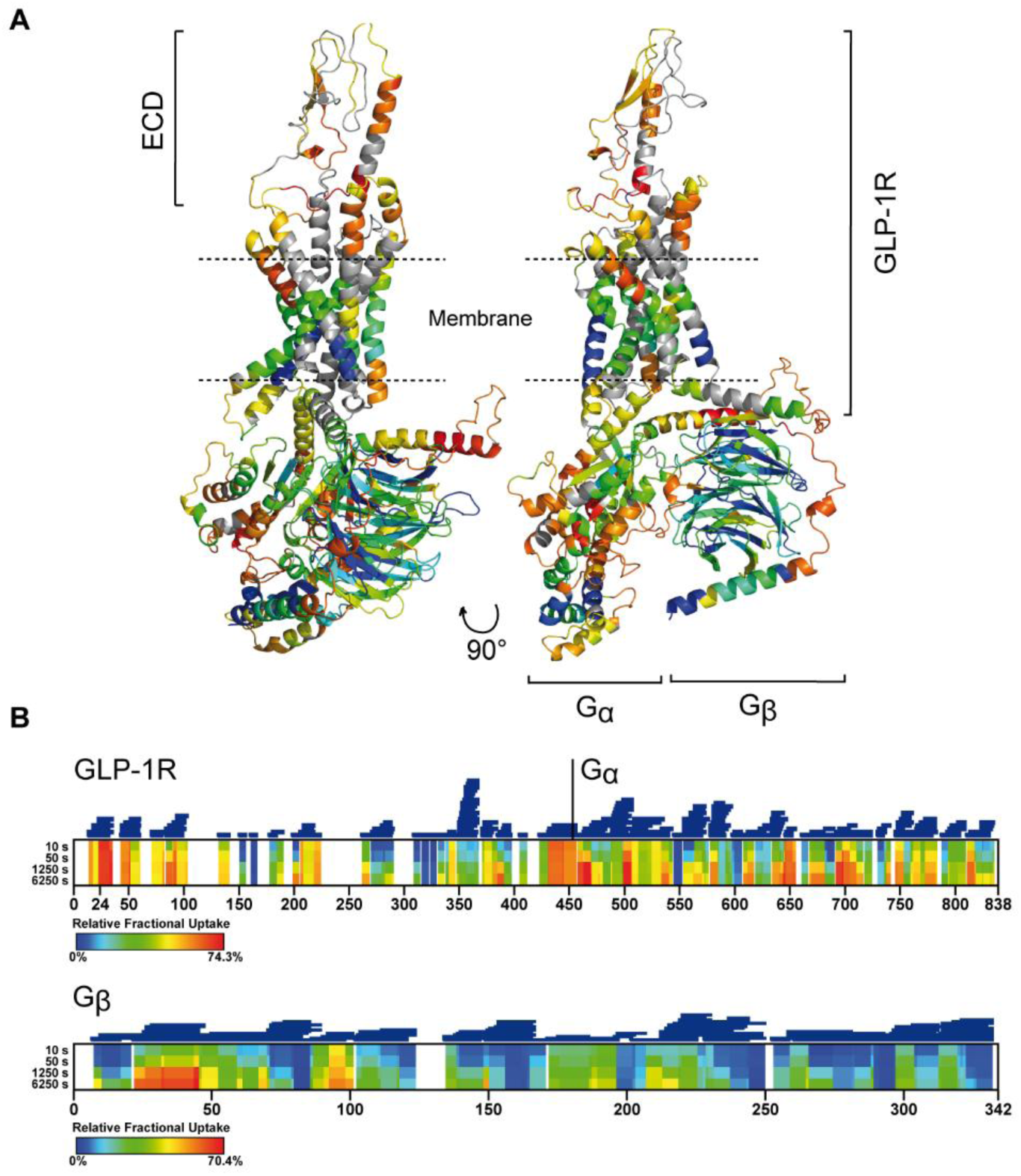
Structural Dynamics of the Pre-Coupled GLP-1R-Gs. HDX-MS results presented as a heatmap using a rainbow scale representing relative from 0% to 70.4%. **(A)** AlphaFold model of the GLP-1R–Gs complex with deuterium incorporation at 6,250 seconds mapped. (**B**) HDX-MS fractional uptake values obtained for the apo GLP-1R–Gs complex (GLP-1R-Gα and Gβ subunit of Gs) over deuteration time points ranging from 10 to 6,250 seconds. Peptide-level resolution is shown, with the positions of the 211 identified peptides indicated by blue bars above the heatmap, corresponding to a sequence coverage of 78.6% and an average redundancy of 3.79. The exclusion of the Gγ subunit reflects the absence of analytical data for this component and enhances the clarity of data visualization. Comparison with an AlphaFold-predicted full receptor-Gs complex revealed no structural differences, particularly within the key interaction interfaces (**Fig. S4**).

HDX-MS analysis of the apo GLP-1R–Gs complex revealed several unstructured loops and flexible helical regions in GLP-1R exhibiting high levels of deuterium exchange (≥50% at t = 10 s), including parts of the ECD, ECL1, ICL3, and ECL3. Apart from ICL3, all of these segments reside in the extracellular portion of GLP-1R and contribute to a critical agonist-binding interface for small-molecule mimetics (28, 29, 47). In contrast, although ICL3 is generally considered to participate in G-protein coupling, its deuteration level reaches a maximum within the first 50 seconds, indicating rapid solvent accessibility and pronounced local disorder. The 7TM of GLP-1R displayed a modest degree of protection over the HDX-MS time course, consistent with its overall compact architecture. The central portions of the TM regions were the most protected from deuteration, likely due to the hydrophobic core of the LMNG/CHS micelle, however, the relatively high levels of deuterium exchange observed for the TM2, TM4, and TM7 termini (≥50% at *t* = 6250 s) are likely attributable to their higher degree dynamical character(s) and resultant solvent exposure. In addition, HDX-MS allowed us to map deuterium-exchange patterns within Gα and Gβ subunits of Gs, revealing areas with time-dependent exchange, characteristic of a folded protein with differences in secondary structure and dynamics.

### GLP-1R-Gs Activation Dynamics with Small-Molecule GLP-1 Agonists

First, we sought to characterize GLP-1R activation dynamics by non-peptide agonists (NPAs) with well-established binding modes and pharmacological profiles, before extending HDX-MS analysis to peptide ligands that have not previously been interrogated in this manner. To accurately design conditions for subsequent HDX-MS experiments on GLP-1R-Gs occupied by NPAs, Chu-128 and danuglipron, their binding was assessed using an affinity selection mass spectrometry (ASMS) assay. High-affinity equilibrium dissociation constants, K_D_ values of 279.5 and 4.63 nM, were determined for the interaction between danuglipron and Chu-128 with our GLP-1R-Gs construct, respectively (**Fig. S5**). These values were employed to guide complex formation for HDX-MS; GLP-1R–Gs was equilibrated with a 10-fold molar excess of either danuglipron or Chu-128 to achieve close to saturation of protein in complex with ligand. We performed HDX labelling at 10, 50, 1,250 (20 min 50 s), and 6,250 seconds (1 h 44 min 10 s) on LMNG-CHS detergent-solubilised GLP-1R-G_s_ in presence or absence of NPA and performed ΔHDX analysis between individual time points, as well as evaluating aggregated differences (∑ΔHDX) to better identify potential peptides that are conformationally active (48–50).

Danuglipron and Chu-128 both induced significant protection to deuterium exchange at the N-terminal regions of the ECD, ECL1, and the extracellular portions of TM1 and TM2 of the GLP-1R helical bundle (**Fig. 3**). Similar protection effects of both agonists propagate deeper into the receptor structure, manifesting as protection of TM4, TM6, and TM7, as well as the intracellular region of TM5 associated with ICL3, and ICL4 linked to helix 8 (**Fig. 3A**, yellow region), suggesting a comprehensive stabilisation of the GLP-1R backbone dynamics as a feature of NPA binding. However, increased deuteration was caused by both NPA within residues 83–103 (83-99 for danuglipron) covering a part of the core ECD structure that forms the binding pocket for the C-terminal portion of GLP-1, suggesting harmonised destabilisation of the pocket region (**Fig. 3A**). Interestingly, differences between danuglipron and Chu-128 were noticeable, with danuglipron causing enhanced stabilisation within the N-terminal of the ECD, TM2 and TM6 compared to Chu-128, with Chu-128 stabilising TM4 more than danuglipron (**Fig. 3A**). Intriguingly, danuglipron, but not Chu-128, stabilised ECL3 connecting TM6 and TM7 domains (**Fig. 3C**, peptide 372-384).

**Figure 3.**
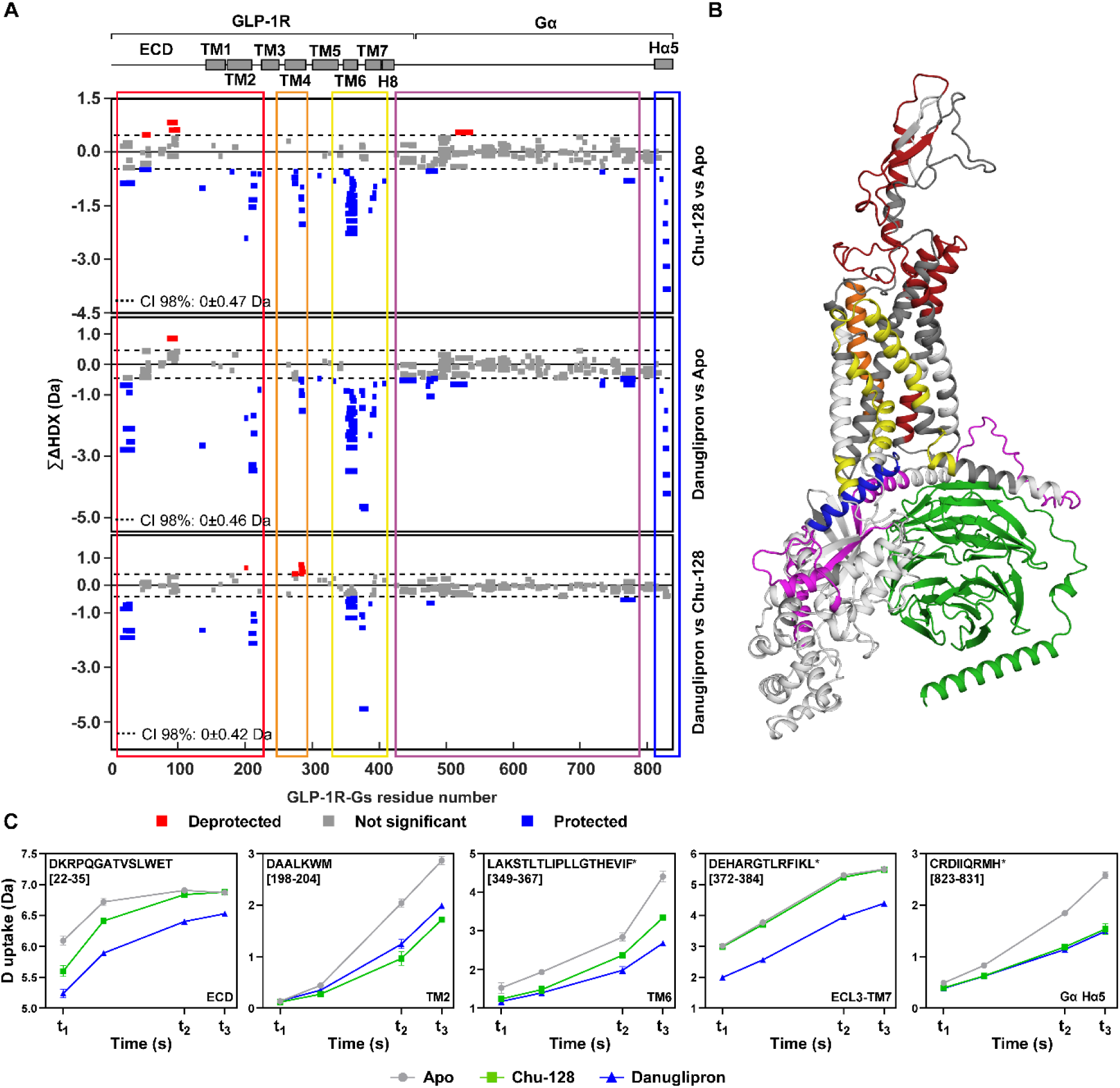
Effect of Chu-128 and danuglipron on GLP-1R-Gs structural dynamics. (A) ∑ΔHDX plots showing two-state comparisons: Chu-128 vs apo, danuglipron vs apo, and danuglipron vs Chu-128. Red signifies peptides with increased HDX between states and blue represents peptides with decreased HDX. Confidence intervals (CI_98%_) are shown as grey dashed lines and grey data are peptides with insignificant ΔHDX. All measurements were performed in triplicate and all supporting HDX-MS peptide data and ΔHDX plots for individual timepoints can be found in the **Source Data file** and **Fig. S6-S8**, respectively. Regions of significant interest are delineated by coloured frames highlighting ECD (red), TM4 (orange), TM6–ECL3–TM7-H8 (yellow), the Gα subunit (purple), and the Gα helix α5 (blue). (**B**) AlphaFold GLP-1R–Gs model pf regions of interest highlighted in (**A**). (**C**) Representative D-label uptake plots for peptides 22-35, 198-204, 349-367, 372-384, and 823-831 and their locations within the GLP-1R–Gs complex. Peptides with identified presence of bimodal distributions are indicated by an asterisk next to each peptide sequence. Uptake plot data are the average with standard deviation from repeated measurements (*n* = 3) plotted but which are often too small to be observed.

Strikingly, the C-terminal helix α5, a critical component of the activated GLP-1R–G_s_ complex, inserting into the cytoplasmic cavity formed by the receptor’s transmembrane helices (TMs) to initiate downstream signalling, becomes significantly protected to exchange by NPA activation (**Fig. 3A-B**, blue region spanning residues 823-835). In the presence of agonist, protection is already evident at 10 seconds and progresses over time, suggesting stabile sequestered stabilisation of helix α5 into receptor’s core (**Fig. 3C**, peptide 823-831). Conversely, the magnitude of this effect on helix α5 does not differ between the two non-peptide agonists tested. Whereas the rest of the Gα and Gβ deuteration profiles were unaltered by NPA binding, there is subtle evidence for modification to the dynamics of regions (469-488 and 760-783) covering the Gα nucleotide-binding catalytic domain of Gs (51, 52) that are specifically modulated by non-peptide GLP-1R agonists (**Fig. 3A-B**, purple region).

### Effect of endogenous peptide ligands on the conformational ensemble of GLP-1R-Gs

Prior to this study, the impact of peptide agonist binding on GLP-1R dynamics was largely unknown, most likely because these amphipathic peptides pose significant experimental challenges due to their adhesive behaviour. In our study, we obtained HDX-MS data for GLP-1R-Gs complex in the presence of GLP-1[7-36] agonist and its inactive metabolite GLP-1[9-36] by enhancing peptide solubility through the addition of the cyclic polypeptide antibiotic bacitracin, which has previously been used successfully to stabilize proteins and peptides in animal cell cultures (53; see **Methods**). Because bacitracin was excluded from the small-molecule HDX-MS experiments, the NPA and peptide ligand ΔHDX datasets are not directly comparable, and any comparisons are interpreted qualitatively.

HDX-MS was performed at 10, 1,250 (20 min 50 s), and 6,250 seconds (1 h 44 min 10 s) for GLP-1[7–36] and GLP-1[9–36] binding experiments. Peptide ligands were added at 2.5-molar excess which, based on the low nanomolar affinity range (1-2 nM) reported in the literature (54), should achieve a dominant peptide-GLP-1R-Gs complex. Even so, minimal deuterium uptake changes were observed across the timepoints; informatively, a previous HDX-MS study on the truncated extracellular domain of GLP-1R (nGLP-1R) also observed insignificant changes to deuterium exchange upon binding GLP-1 in comparison to NPAs, this trait may indicate that GLP-1 peptide ligand binding interactions are primarily mediated through side-chain interactions, or that any stabilization to amide hydrogen is close to limit of detection (74). However, an ∑ΔHDX analysis revealed significant areas of deuterium uptake change, making it possible to locate areas with altered structural dynamic profiles (**Fig. 4A**).

**Figure 4.**
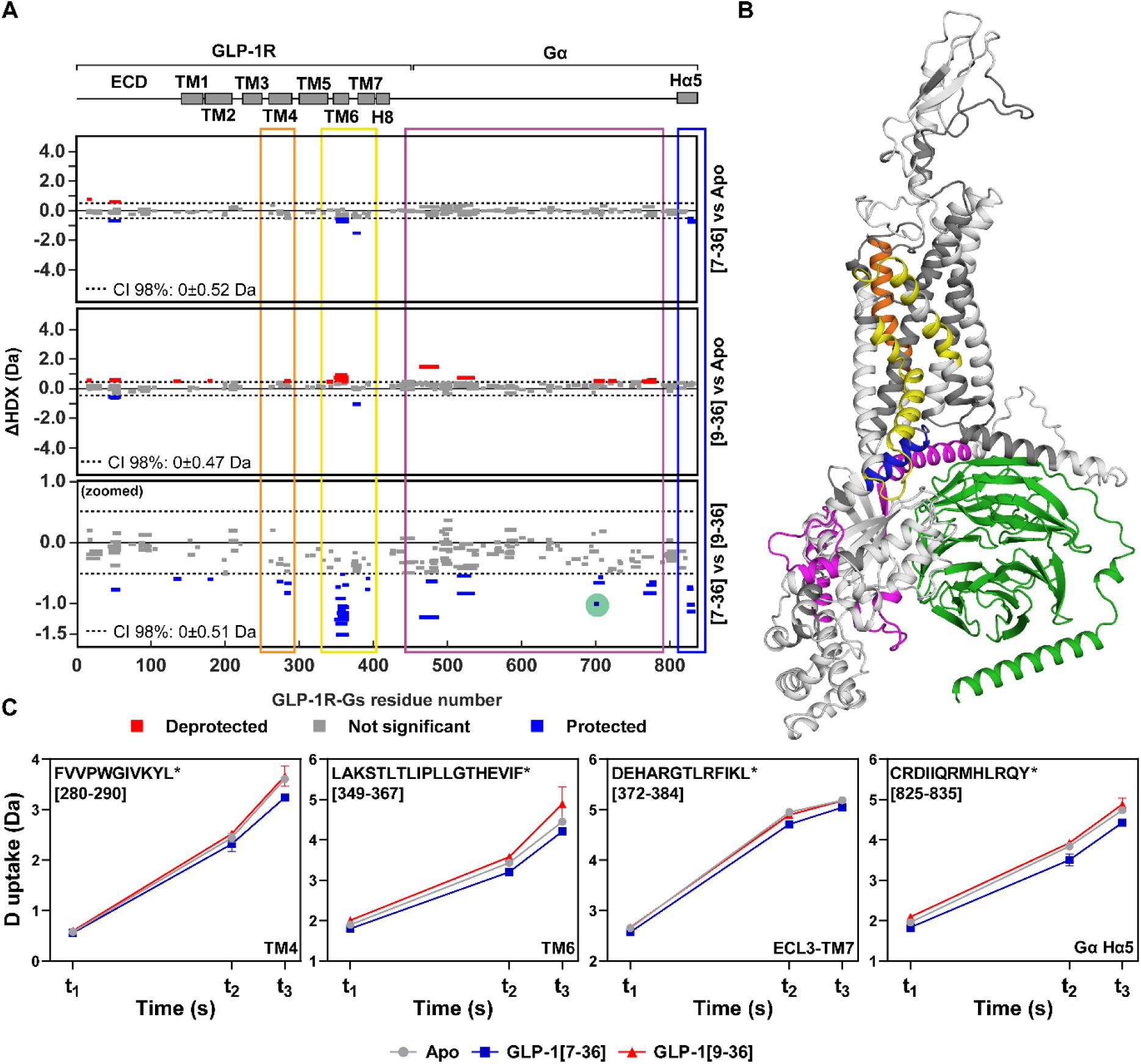
Distinct effects of GLP-1[7-36] and GLP-1[9-36] on GLP-1R-Gs structural dynamics revealed by ∑ΔHDX statistical analysis. ∑ΔHDX plots showing two-state comparisons: GLP-1[7–36] vs apo, GLP-1[9–36] vs apo, and GLP-1[7–36] vs GLP-1[9–36] (**A**). Red signifies peptides with increased HDX between states and blue represents peptides with decreased HDX. Confidence intervals (CI_98%_) are shown as grey dashed lines and grey data are peptides with insignificant ΔHDX. Peptide 698–704 is not included in the first two graphs for clarity due to its high deuterium uptake; however, it is highlighted by a light green circle in the bottom plot for comparison of GLP-1[7–36] vs GLP-1[9–36]. All measurements were performed in triplicate and all supporting HDX-MS peptide data and ΔHDX plots for individual timepoints can be found in the **Source Data file** and **Fig. S9-S11**, respectively. Regions of significant interest are delineated by coloured frames highlighting TM4 (orange), TM6-ECL3-TM7 (yellow), the Gα subunit (purple), and the Gα helix α5 (blue). The AlphaFold GLP-1R–Gs structure highlighting regions of interest is shown in (**B**). Panel (**C**) depicts peptides 280–289, 349–367, 372–384, and 823–835 and their locations within the GLP-1R–Gs complex. The presence of bimodal distributions in peptide spectra is indicated by an asterisk next to each peptide sequence. Uptake plot data are the average and standard deviation from repeated measurements (*n* = 3).

Regarding ECD binding, both ligands showed mixed exchange effects in the 45–60 region (**Fig. 4A**, GLP-1[9–36] vs Apo). Interestingly, two arginine residues, R43 and R44, showed protection to exchange in both peptide-bound states (**Fig. 4A**). Most significant, was the diametrically opposed effects uncovered between GLP-1[7-36] and GLP-1[9-36] on areas of ECD-TM1, ECL3-TM7 and, most significantly, within TM6 regions of GLP-1R; where GLP- 1[7-36] binding causes decreased deuterium uptake but GLP-1[9-36] increased exchange. Moreover, GLP-1[7-36] caused deuterium protection within the Gα C-terminus, whereas GLP- 1[9-36] did not. Collectively, these findings suggest that the GLP-1[9–36] metabolite promotes increased dynamics within the TM6–ECL–TM7 region while fails to stabilise the Cα-terminus, which contrasts with the stabilising effects observed for GLP-1[7–36] and NPAs and may define its inactive classification.

### Bimodal HDX-MS spectra suggest interconversion between conformations within the conformational ensemble of GLP-1R-Gs

In HDX-MS studies of membrane proteins, particularly transporters, regions that display slow, coordinated unfolding/refolding motions consistent with bimodal spectra (40) are usually those whose dynamics are influenced by substrate binding (55, 56). Interestingly, in our study, several regions of the pre-coupled GLP-1R–Gs complex in the apo state exhibited bimodal isotopic envelopes with distinct low- and high-mass populations whose intensities interconverted either partially or fully over the experimental time. In GLP-1R, slow unfolding/refolding occurs in region 270-290, restricted to the intracellular portion of TM4, and 349-367 spanning the entire TM6, yet only TM6 conformational interconversion has been found to be significantly modulated by non-peptide agonist binding (**Fig. 5A**). While the two populations continue to interconvert in intensity after 1,250 seconds of exchange, NPAs prolong their lifetime, as indicated by relative distribution between the low-mass (closed, unlabelled) and high-mass (open, labelled) states remaining unchanged over the time (**Fig. 5C**). In contrast, no stabilising effect of NPAs on TM4 was observed.

**Figure 5.**
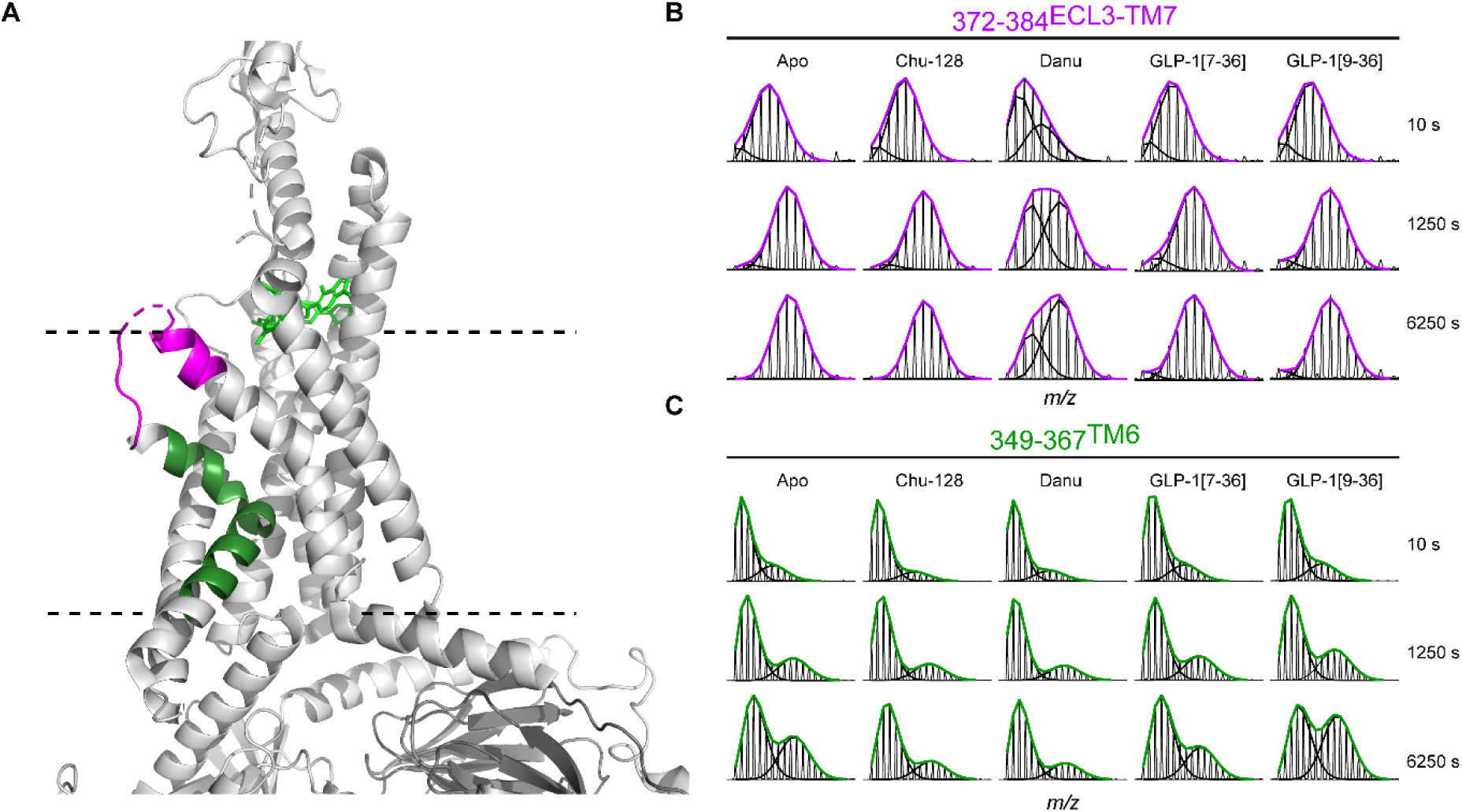
Allosteric modulation of the GLP-1 receptor by peptide and non-peptide ligands. (**A**) Structure of GLP-1R (PDB:6X19) in complex with Gs and Chu-128, with TM6 and ECL3-TM7 regions highlighted in green and purple, respectively. Complex is positioned relative to the membrane, depicted by two dashed lines, with the Orforglipron moiety shown in green. HDX-MS plots show bimodal distributions in GLP-1R ECL3–TM7 (residues 372–384; **B**) and TM6 (residues 349–367; **C**), indicative of conformational heterogeneity. Spectra were analysed using HX-Express (v3) and further edited in Prism 9 (version 9.5.1.733). Distinct populations are shown as black lines, and mixed envelopes as lines coloured according to region.

Beyond that, bimodal spectral signatures were found in multiple peptides for region 372-396 spanning ECL3 and TM7, which we defined as a danuglipron-sensitive region in the extracellular portion of the receptor (**Fig. 3A**, yellow region; **Fig. 3C**, peptide 372-384). However, peptide 372-384 included in this region was found to exhibit spectral bimodality not only in danuglipron-bound state, but also, more subtly, in Chu-128-bound and Apo states (**Fig. 5B**). Notably, danuglipron markedly reduces the interconversion rate, thereby stabilizing the low-mass (closed, unlabelled) state. In contrast, both the apo form and the Chu-128–bound state is dominated by a fast-exchanging population since the initial time point. We propose that danuglipron, but not Chu-128, restricts conversion of long-lasting local conformations within ECL3 and extracellular portion of TM7, which highlights differences between the NPAs examined and which is tempting to link to the full and partial agonism of danuglipron and Chu-128, respectively. In GLP-1 peptide-bound states, we also observed bimodal distribution within parts of the TM4, TM6 and ECL3-TM7 of GLP-1R. Like NPAs GLP-1[7–36] does not alter the lifetimes of these populations in the TM4 region relative to the apo state, whereas GLP-1[9–36] facilitates their interconversion (**Fig. S12**, peptide 280-288). By contrast, like NPAs, GLP-1[7–36] prolongs the lifetimes of these populations in TM6 (peptide 349-367), reducing interconversion, while GLP-1[9–36] steadily promotes their conversion. Notably, peptide agonists induced protection of region ECL3-TM7 (peptide 372-384) but did not stabilise slow- and fast-exchanging populations in the manner observed for danuglipron (**Fig. 5B**).

Furthermore, localised regions with bimodal spectral features were also found in Gα C-terminus spanning residues 823-835, where rapid conformational conversion around 1250 seconds was observed, a process that is delayed in the presence of NPAs (**Fig. 6A-B**). This suggests that NPAs favour a closed, backbone protected helix α5 conformer, probably caused by a tighter sequestration upon ligand binding. This is perhaps a defining allosteric effect of NPAs, possibly allowing their particular GDP/GTP nucleotide exchange events to occur. Moreover, we found that peptide agonists, but not danuglipron or Chu-128, uniquely induced bimodality of peptide 698-704 which is absent in the apo state (**Fig. 6A** and **C**). This sequence region represents the catalytic Gs protein switch III loop, which is unresolved in both GLP-1-bound and non-peptide agonist-bound structures, likely due to its high flexibility (28, 29). The presence of two distinct populations that remain unchanged over the time course likely reflects no, or very slow, GLP- 1 (incretin) induced existence of, putatively, partially and fully opening switch III states.

**Figure 6.**
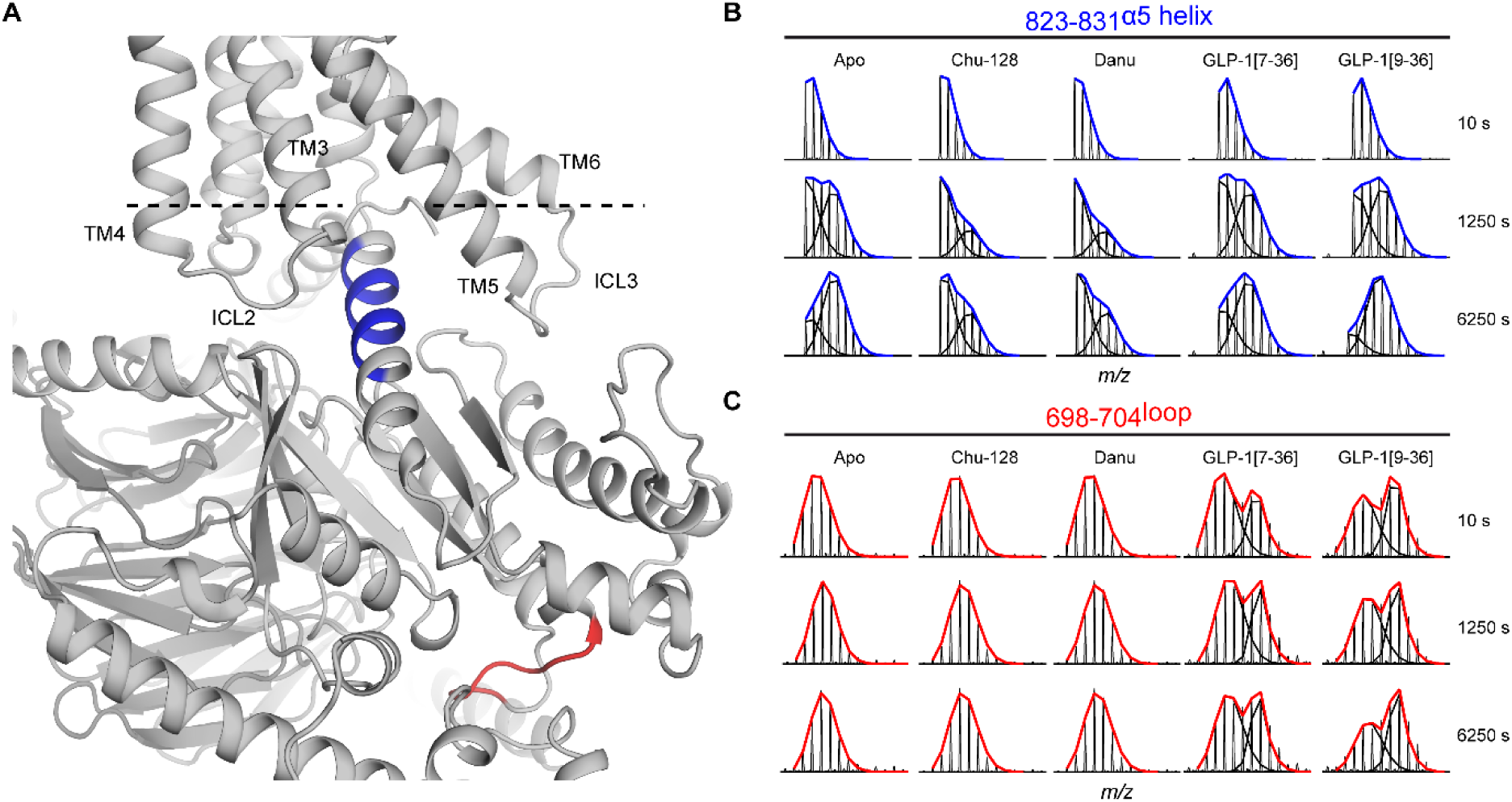
Allosteric modulation of the α5 helix and catalytic domain of Gα subunit by peptide and non-peptide ligands. (**A**) AlphaFold structure of GLP-1R-Gs complex, with 698-704 and 823-831 regions of Gα subunit highlighted in red and blue, respectively. The complex is positioned relative to the membrane, with a dashed line delineating the boundary between the transmembrane and intracellular areas. HDX-MS plots showing bimodal distribution in Gα subunit α5 helix (residues 823-831, **B**) and catalytic domain (698-704, **C**), an indication of conformational heterogeneity. Spectra were analysed using HX-Express (v3) and further edited in Prism 9 (version 9.5.1.733). Distinct populations are shown as black lines, and mixed envelopes as lines coloured according to region. The heterogeneity in region α5 helix is modulated by ligands. The heterogeneity in region of 698-704 is induced solely by addition of peptide ligands and modulated differently.

## Discussion

It is well-known that ligand-bound GLP-1R activates G proteins to drive downstream cellular responses (6), but the molecular mechanism underlying this process remains unclear. Here we present structural insights into the dynamics of pre-coupled GLP-1R-Gs state using HDX-MS. Comparing ligand-evoked similarities and differences across GLP-1R-Gs structural dynamics between established GLP-1 mimetic (non-peptide agonist, NPA) drugs allowed us to define consensus and defining protein regions modulated by NPA activation. Simultaneously, the inclusion of both the endogenous agonist GLP-1[7-36] and the inactive metabolite GLP-1[9- 36] highlighted structural and dynamic signatures between natural GLP-1 ligands and GLP-1 peptide mimetics, as well as potentially differentiate the structural dynamics that underpin agonism from antagonism. For example, NPAs examined in our study show substantial engagement with the ECD, consistent with danuglipron adopting a more closed conformation and Chu-128 a more open conformation relative to each other (28). In contrast, peptide ligand did not provide significant evidence of the same degree of ECD engagement, likely because this interaction is manifested via residual side-chain interactions rather than triggering significant conformational modulation. Taken together, our HDX-MS analysis demonstrates that while agonists induce broadly similar conformational effects on the GLP-1R, the magnitude of these changes together with the allosteric signals propagated to the G protein differ substantially. These differences are likely rooted in distinct ligand binding modes and provide a mechanistic basis for the divergent signalling profiles reported for Chu-128, danuglipron, and GLP-1[7–36].

### GLP-1[9-36], but not Chu-128, danuglipron, or GLP-1[7-36], conformationally restricts TM4

Previously, no direct engagement of TM4 in GLP-1R activation has been proposed, as no direct ligand–TM4 interactions have been observed (29). TM4 is generally considered a rigid peripheral transmembrane helix exhibiting minimal positional changes during activation of many GPCRs. However, TM4 involvement in receptor activation has been reported for class F GPCRs (57). In class A GPCRs, subtle TM4 movements have also been proposed to be essential for signal transduction from the ligand-binding pocket to the G-protein interface, thereby contributing to receptor G-protein coupling specificity (58). In GLP-1R, however, only modest TM4 rearrangements have been observed, restricted to the helix termini, with no detectable displacement within the membrane-embedded region of TM4 (28, 57, 59).

In our HDX-MS analysis, we observed significant protection to exchange for TM4 upon binding of the NPAs tested, with a stronger effect for Chu-128 versus danuglipron (**Fig. 3A**, orange). While this region exhibits greater protection to exchange in the GLP-1[7–36]-bound state when compared to a GLP-1[9–36]–bound state (**Fig. 4A**, [7-36] vs [9–36]). HDX-MS also revealed that regions within the TM4 exhibit bimodal distribution in apo state and all four ligand-bound states examined, yet only GLP-1[9-36] facilitates interconversion between these populations (**Fig. S12**, peptide 280-288).

We propose that the observed protection of TM4, existing in two distinct populations over the experimental timescale, remains compatible with GLP-1R activation. We hypothesize that this behaviour reflects differing packing of TM4 against TM3 shifting its transmembrane helical backbone stability within a pre-coupled complex and during receptor activation, rather than TM4 experiencing extensive rigid-body structural rearrangements (comparable to TM6) not evidenced in ligand-bound high-resolution structural models. Although TM4-TM3 contacts have not been previously reported for GLP-1R, this hypothesis is supported by HDX-MS data showing altered dynamics in TM4 region encompassing aromatic residues Y269, W274, F280, and W284, along with I272 and V281, whose orientations are favourable for stabilizing hydrophobic interactions with TM3. Considering this interpretation, the favouring of long-lived TM4 backbone opening observed in the GLP-1[9–36]–bound state likely reflects receptor deactivation, manifested by a shift toward a less stable complex characterised by weakened TM4-TM3 hydrophobic interactions. We propose that mapping the dynamics of the TM4 domain could be valuable for detailed drug screening, as it represents an HDX feature that clearly distinguishes agonists from antagonists in their ability to stabilize the fully active receptor state.

### Ligands with differing modes of binding restricts TM6 and helix-α5 movement

In G-protein coupled and ligand-bound states, GLP-1R shows outward movement of TM6 with introduced kink within its middle section likely in order to accommodate helix α5 of GαGs subunit within receptor activation (44). This event breaks the tight TM6–TM3 coupling, and partially contacts with TM5 and TM7, a hallmark of class B GPCR activation (60). This all together promotes formation of the pre-coupled GPCR–Gs complex thought to occur *in vivo* (61). Because GLP-1R and Gs were examined in a pre-coupled state, we did not assess dynamic changes accompanying complex formation. Here, our HDX-MS analysis identified TM6 and the Gα α5 helix as ligand-responsive regions.

TM6 dynamics are particularly noteworthy, as they refine and extend conclusions drawn from static structural studies of GLP-1R activation (62–64). Specifically, our HDX-MS results demonstrate that agonists only promoted TM6 protection to exchange, consistent with TM6 backbone stabilisation for achieving receptor activation, together with tight sequestration of the Gα α5 helix. According to our findings, agonist binding seems to promote formation of a fully activated GLP-1R-Gα-GDP complex by driving deeper α5-helix insertion and stabilisation, which could subsequently facilitate opening of the GDP-binding clefts of Gα to enable nucleotide exchange, the event necessary for cellular response (61). With corresponding dynamic changes in the Gα α5 helix symbolising differences in G-protein coupling efficacy.

Additional regions whose structural dynamics perturbations may reflect enhanced G-protein coupling include the intracellular portion of TM5 and the region spanning ICL4 and helix 8, both of had areas which were significantly protected to exchange in danuglipron- and Chu-128-bound states. In contrast, the TM5–ICL3 region was deprotected to exchange in the GLP-1[9-36]-bound state, further underscoring the distinct dynamic signatures associated with agonism and antagonism at GLP-1R. This together with concerted deprotection of TM4, TM6, and the Gα α5 helix, along with enhanced interconversion dynamics, supports the conclusion that GLP-1[9–36] actively induce destabilisation of the active GLP-1R-Gs conformation.

### ECL3-TM7 modulation defines Chu-128 partial agonism

The ECL3 was previously considered a short peptide linker that merely connects the functionally important transmembrane domains TM6 and TM7. More recently, however, ECL3 has attracted increasing interest due to its emerging role in GPCR signalling, including agonist binding, ligand selectivity, and receptor activation (65). The significance of ECL3 stems from its position linking TM6 and TM7, whose cytoplasmic ends are connected to ICL3 and helix 8, respectively, domains that are directly implicated in GPCR activation and downstream second-messenger production (66). As a result of its placement, ECL3 is thought to regulate receptor activation by constraining the relative movements of TM6 and TM7, with activation involving reciprocal release or rearrangement of these constraints. This concept is supported by activating mutations within the ECL3 of the δ-opioid receptor, which relieve such structural restraints and promote receptor activation (67). Additional evidence comes from studies of chimeric β2-adrenergic receptors in which ECL3 was substituted with that of either the α_1a_-adrenergic (α_1a_AR) or thyrotropin receptors (68, 69). Substitution with α_1a_AR ECL3 produced a constitutively active chimeric receptor with increased basal activity, likely due to relaxed conformational constraints on TM6 and TM7 relative to the wild-type receptor. In contrast, substitution with thyrotropin receptor ECL3 resulted in a reduced thyrotropin-stimulated cAMP response. These constraints may arise from interactions between ECL3 and the receptor’s extracellular domain (ECD), mediated through hydrogen bonding or disulfide bond formation (70, 71). Like class A GPCRs, the class B GLP-1R has also showed evidence of ligand-induced reorganisation of the TM6-ECL3–TM7 region, observed upon binding of both peptide and non-peptide agonists (47, 72, 73). Together, these findings support the hypothesis that ECL3 is not merely a passive linker, but a critical origin point for the GPCR activation pathways.

In this context, our HDX-MS study reveals ligand-evoked structural dynamics reorganisation of the TM6–ECL3–TM7 region to be a key driver of agonism extent. This is evidenced by distinct exchange-competence differences between full agonist (danuglipron-bound) and partial agonist (Chu-128-bound) states. Chu-128 did not show dynamical change of ECL3 whereas danuglipron, uniquely, stabilised an otherwise cooperative interconversion between backbone closed/protected and open/deprotected populations in ECL3-TM7. Unexpectedly, these findings differ from previous conclusions which suggested that danuglipron closely mimics GLP-1 peptide activation modes (28), leading us to propose that ECL3 perturbation is a defining feature of danuglipron-mediated full agonism. This engagement likely underlies the more pronounced TM6 rearrangements induced by danuglipron compared with Chu-128, as observed by HDX-MS (**Fig. 3A**, danuglipron vs Chu-128). Taken together, these differences in dynamics are consistent with distinct signalling profiles of these two non-peptide agonists, namely full versus partial agonism.

### Only GLP-1 peptides selectively modulate catalytic domain of Gsα

Upon GPCR activation, the Gsα protein undergoes GDP–GTP exchange, triggering major conformational rearrangements in its switch regions, including switch III. Switch III is known to play a central role in effector activation by mediating interactions between the Gα subunit and adenylyl cyclase, thereby promoting cAMP production. Binding of GTP stabilises switch III, along with switches I and II, into an active conformation that facilitates dissociation of the Gα subunit from the Gβγ dimer and sustains downstream signalling. GLP-1-induced spectral bimodality was observed for peptide 698-704 spanning portion of catalytic domain of GαGs, indicating triggering of local, substrate-dependent unfolding intermediate switch III states which may reflect rearrangements in the catalytic domain of Gα associated with its transition into a nucleotide-exchange-competent state, leading to G protein activation. Notably, this was observed for GLP-1 peptides exclusively, and not for small-molecule NPAs, suggesting it could be functioning as an allosteric switch that primes Gα for nucleotide exchange for peptide-driven GLP-1R activation specifically.

### Limitations of the study

It should be emphasised that HDX-MS probes protein backbone dynamics rather than side-chain interactions directly. As a result, GPCR microswitch activation events that produce only subtle or localised structural changes may remain undetected or fall below the method’s sensitivity threshold (75). Importantly, the comparative analysis of apo and holo state dynamics required a stable ligand-free GLP-1R:Gs complex, which was achieved by covalently fusing GLP-1R to a nucleotide-free favoured Gα subunit (achieved through mutation of the GTP binding site) while maintaining the association of the Gβ and Gγ subunits. Although this construct was critical to achieving differential HDX-MS it is an engineered construct which may not recapitulate the entire *in vivo* conformational ensemble. However, a notable advantage of this approach is the avoidance of receptor thermostabilising mutations enabling ligand-independent purification without artificial stabilisers like nanobodies or positive modulators (29, 76, 77). Moreover, GLP-1R-Gs was investigated within the “gold-standard” LMNG/CHS cholesterol-containing detergent micelle membrane mimetic which is known to produce biologically relevant GPCR states for structural biology investigation. However, it is devoid of the endogenous lipids that have been observed to be important for modulating GPCR-G protein signalling (78–80). Future studies could implement emerging HDX-MS strategies (mainly developed on bacterial membrane systems to date) which mitigate the analytical challenges on studying membrane protein-lipid assemblies to achieve insight into structural dynamics of GLP-1R within a natural lipid surround (81). This would provide potentially more relevant conformational ensembles, enabling a more refined understanding of functional dynamics.

## Materials and Methods

### Purification of GLP-1R:Gs non-covalent complex

40 mL of membrane fraction containing co-expressed GLP-1R:Gs complex, obtained from 20 L of culture, were ultracentrifuged at 100,000 × g for 1 h at 4 °C to remove storage buffer. The pellet was resuspended in 30 mL of pre-chilled buffer A (150 mM NaCl, 30 mM HEPES, 8 mM MgCl_2_, 5 mM CaCl_2_, pH 7.5) supplemented with protease inhibitor tablet (Roche), PMSF (Thermo), and apyrase (100 µL of 10 mg/mL stock (3.0 U/mg); Sigma-Aldrich), and incubated on ice for 30 min. Detergents were then added to final concentrations of 0.5% LMNG and 0.05% CHS from stock of buffer A containing 4% and 0.4% of LMNG and CHS, respectively, and the volume was adjusted to 50 mL with buffer A. The mixture was incubated for an additional 30 min for solubilisation. Solubilised material was incubated overnight at 4 °C with gentle mixing with 1 mL of PureCube Co-NTA agarose (Cube Biotech) pre-equilibrated in buffer A containing 0.5% LMNG and 0.05% CHS for IMAC purification.

The following day, Co-NTA agarose was washed with 5 mL buffer B (150 mM NaCl, 30 mM HEPES, pH 7.5, 0.01% LMNG, 0.001% CHS), followed by two washes with 5 mL buffer B supplemented with 50 mM imidazole. Bound GLP-1R:Gs was eluted with 2 × 1.25 mL of buffer B containing 150 mM imidazole, with a 5-min incubation for each volume prior to elution. Elution fractions were pooled, buffer-exchanged into buffer C (150 mM NaCl, 30 mM HEPES, pH 7.5, 0.01% LMNG, 0.001% CHS), concentrated using a 100 kDa MWCO centrifugal filter (Amicon), and flash-frozen prior to size-exclusion chromatography (SEC).

For SEC, a 475 µL aliquot was thawed on ice, supplemented with TCEP to a final concentration of 100 µM, and loaded onto a Superose™ 6 Increase 10/300 GL column (Cytiva) equilibrated in buffer D (150 mM NaCl, 30 mM HEPES, pH 7.5, 0.01% LMNG, 0.001% CHS, 100 µM TCEP). Chromatography was performed at 0.5 mL/min using an AKTA Pure FPLC system. Fractions 12–27 corresponding to GLP-1R:G*_s_* were collected, analysed by SDS–PAGE, and GLP-1R and G*_s_*γ (6×His-tagged proteins) were detected by immunoblotting using an anti-6×His monoclonal antibody conjugated to horseradish peroxidase (HRP; 1:10,000 dilution). Fraction 20 was subjected to a second SEC under identical conditions, except for a flow rate of 0.3 mL/min. Fractions 19–22 from the single peak were analysed as above, and fraction 19, showing highest purity and sufficient GLP-1R signal (50–60 kDa), was selected for HDX-MS. Tandem SEC results are shown in Fig. S1A–D in *SI Appendix*.

### Purification of the apo-active GLP-1R–Gαs–Gβγ complex

The GLP-1R–Gαs fusion construct encodes human GLP-1R (residues 24–452) with an engineered Gαs subunit appended directly to the receptor C-terminus. An N-terminal CD8α signal peptide followed by a FLAG tag precedes GLP-1R residue 24. The Gαs component comprises human Gαs long isoform residues 26–394 bearing mutations favoring the nucleotide-free state (S54N, G226A, E268A, N271K, K274D, R280K, T284D, I285T, and A366S), with its native N-terminus (residues 1–25) replaced by the corresponding Gαi1 N- terminal sequence (residues 2–18: GCTLSAEDKAAVERSKM) to facilitate Gβγ heterotrimer assembly. The Gβ1 subunit comprises canonical residues 2–340 with an N-terminal MAL cloning artifact. The Gγ2 subunit comprises canonical residues 2–70 preceded by an N-terminal 8×His tag. Full amino acid sequences are provided below.

Sf9 cell pellets co-expressing the GLP-1R–Gαs fusion, Gβ1, and Gγ2 from recombinant baculoviruses were resuspended in lysis buffer containing 10 mM HEPES (pH 7.5), 50 mM NaCl, 5 mM CaCl_2_, 2 mM MgCl_2_, Turbonuclease (Accelagen), apyrase (NEB), and protease inhibitors (Thermo Fisher Scientific). The suspension was incubated for 1 h at room temperature, during which Turbonuclease degraded genomic DNA and RNA to reduce lysate viscosity and apyrase hydrolyzed GDP and GTP to promote complex formation in the nucleotide-free state. Membrane proteins were solubilized by addition of lauryl maltose neopentyl glycol (LMNG) and cholesteryl hemisuccinate (CHS; Anatrace) to final concentrations of 0.5% and 0.05% (w/v), respectively, with gentle mixing for 1 h at 4 °C. Insoluble material was removed by ultracentrifugation at 200,000 × *g* for 30 min at 4 °C, and the clarified supernatant was applied to 2 mL bed volume anti-FLAG M2 agarose resin (Sigma-Aldrich) and incubated overnight at 4 °C with gentle agitation.

The resin was transferred to a gravity-flow column and washed with 20 column volumes of wash buffer (10 mM HEPES pH 7.5, 100 mM NaCl, 0.01% LMNG, 0.001% CHS). Bound complex was eluted in the same buffer supplemented with 0.1 mg/mL FLAG peptide. The eluate was concentrated using a 100 kDa MWCO centrifugal concentrator (Amicon) prior to size-exclusion chromatography (SEC).

As a first SEC polishing step, the concentrated eluate was injected onto a Superose 6 Increase 10/300 GL column (Cytiva) pre-equilibrated in SEC buffer (10 mM HEPES pH 7.5, 100 mM NaCl, 0.01% LMNG, 0.001% CHS). Fractions corresponding to the monodisperse complex peak were pooled and concentrated, then subjected to a second SEC step on a Superdex 200 Increase 10/300 GL column equilibrated in the same buffer to remove residual aggregates and lower-molecular-weight contaminants. The final monodisperse complex was concentrated to 3 mg/mL (∼20 µM) for hydrogen–deuterium exchange mass spectrometry (HDX-MS) analysis.

### Cyclic AMP accumulation assays

Recombinant cDNAs encoding human GLP-1R-Gαs fusion proteins that lack the N-terminal FLAG tag were cloned into high-expression pcDNA3.1 vectors. Parental HEK293 cells were adherently passaged in growth medium, transfected in suspension with Promega Fugene6 at a 6:1 reagent to plasmid DNA ratio in growth medium lacking antibiotics, and allowed to adhere to tissue culture flasks in a humidified 37 °C, 5% CO2 environment. Following approximately 48hr of propagation, cells were quickly lifted enzymatically with Gibco TrypLE and cryopreserved with controlled rate freezing and 10% DMSO as a cryoprotectant. Ligands were formulated in DMSO and frozen in aliquots. Concentration response curves were created with direct dilution using a Beckman Coulter Echo 655 acoustic liquid handler that dispensed into Corning 3570 white microtiter plates containing cell assay medium (Dulbecco’s modified eagle medium containing high glucose Gibco 31053, 1X Gibco GlutaMAX, 20 mM HEPES pH 7.5, 0.1% bovine casein Millipore Sigma C4765, 500 μM phosphodiesterase inhibitor 3-isobutyl-1- methylxanthine (IBMX) Millipore Sigma I7018). On the day of the assay, a single assay-ready vial of cells was rapidly thawed to measure intracellular cAMP accumulation. Freezing medium was exchanged with cell assay medium lacking IBMX. Cells were counted for viability and allowed to quiesce for 1hr at 37C following thaw. The suspended cells were added to prewarmed compound treatment plates at equal volume using a ThermoFisher Multidrop Combi dispenser. Final DMSO concentration was 1.1%. Duration of treatment was 30 minutes at 37C. Cyclic AMP was quantified with homogenous time-resolved fluorescence generated from a BMG Labtech Pherastar FSX multilabel reader (337nm excitation) and the Revvity 62AM4PEJ Gs Dynamic Assay with manufacturer’s two step detection protocol. Data were analyzed by the 665nm/620nm emission ratio method, calibrated to external standards in a parallel processed plate, and reported as percent activation compared to native GLP-1(7–36) peptide. Percent values were fit to the four-parameter logistic with variable slope model in GraphPad Prism 10 software. Representative response curves represent three independent run dates.

### Radioligand binding

Radioligand binding was performed as previously described (82), with some slight modification in the buffer and using antibody capture where described below. Sf9 membrane (0.5 - 2.77 µg) expressing the FLAG-GLP-1R-Gs fusion + βγ and 0.1 mg wheatgerm agglutinin polyvinyl toluene scintillation proximity assay beads (WGA-PVT SPA beads, Revvity Cat# RPNQ0001) were incubated with a concentration-response curve of compound or peptide and [^125^I]Tyr^19^- GLP-1(7–36)NH_2_ (∼0.1 nM final, Revvity, Waltham MA) in 1.0 mM CaCl_2_ (Sigma Cat# 21115), 2.5 mM MgCl_2_ (Boston BioProducts Cat# BM670), 0.003% (v/v) Tween-20 (Roche Cat 11332465001), 0.1% (w/v) Bacitracin (Thermo Scientific Chemicals Cat# J62432-14), 25 mM HEPES (Gibco Cat# 15630-080) pH to 7.4 with KOH, in a 200 µL assay volume. The assay was incubated overnight at 25 °C, then centrifuged at 1000 rpm (Beckman Avanti J-15R). Radioligand bound to the GLP-1R expressing membrane (i.e., in close proximity to the WGA- PVT SPA bead) was detected using a PerkinElmer 2450 MicroPlate Counter.

Radioligand binding to purified protein (1 pmol) FLAG-GLP-1R-Gs fusion + βγ was performed as described above with the addition of 0.01% LMNG (laurel maltose neopentyl glycol) and 0.001% CHS (cholesterol hemisuccinate, tris salt) to the assay buffer and was detected by antibody capture using 0.5 mg Anti-Mouse IgG PVT antibody binding beads (Revvity RPNQ- 0017) in place of WGA-PVT SPA beads with the addition of 2.5 µg/mL (final concentration) Monoclonal Anti-FLAG M2 IgG1 (Sigma Aldrich Cat# F3165).

Affinity (K_d_) and expression density (B_max_) values were obtained from homologous competition analysis of human [^127^I]Tyr^19^-GLP-1(7–36)NH_2_ (CPC Scientific, San Jose, CA) versus human [^125^I]Tyr^19^-GLP-1(7–36)NH_2_ using the competitive binding equation one site homologous (GraphPad Prism version 10.6.1 for Windows, GraphPad Software, Boston, MA USA, www.graphpad.com).

### Affinity Selection Mass Spectrometry (ASMS) assay

ASMS binding assays were conducted using a filter-plate–based approach. Saturation binding studies were performed in triplicate in a final volume of 200 µL containing 10 mM HEPES (pH 7.5), 100 mM NaCl, 2 mM MgCl₂, 0.01% LMNG, and 0.001% CHS. Ligands (Chu-128 and danuglipron) were serially diluted to 4× the final incubation concentrations. Each incubation mixture comprised 50 µL ligand solution, 50 µL assay buffer, and 100 µL purified GLP-1R protein (2 µg), followed by incubation for 1 h at 25 °C. Non-specific binding was assessed by substituting assay buffer with 50 µL Orforglipron (2 µM), yielding a final concentration of 500 nM. After incubation, samples were vacuum-filtered through 1 µm glass fiber filters (pre-soaked in 0.25% PEI for 30 min at 4 °C, rinsed, and prewashed with cold assay buffer). Filters were washed five times with 200 µL ice-cold assay buffer, then dried at 55 °C for 3 h. Calibration curves for LC-MS analysis were prepared in eluent (50% methanol, 50% 5 mM ammonium formate) both with and without 500 nM Orforglipron. Following drying, 200 µL eluent was added to each well, and flow through was collected for LC-MS analysis using an MRM method targeting ligand-specific fragments. Calibration curves were applied to quantify ligand concentrations based on signal intensity. Saturation binding data were analyzed using one-site total, non-specific, and specific binding models in GraphPad Prism.

### Preparation of ligands for hydrogen/deuterium mass spectrometry

Danuglipron (LSN3535219) and Chu-128 (LSN3955529) possess high affinity binding to GLP-1R-Gs within LMNG/CHS detergent micelles (Equilibrium dissociation constants (K*_D_*) of 4.63 and 279.5 nM for Chu-128 and danuglipron, respectively), as measured by ASMS (**Fig. S5**). The compounds are soluble in organic solvents and tolerate mixing with aqueous solutions, but require an intermediate dilution step to minimize aggregation during mixing. Therefore, initial stock solutions of danuglipron and Chu-128 at concentrations of 18 mM and 26 mM, respectively, were prepared by dissolving the lyophilized compounds (Eli Lilly and Co.) in DMSO. Small aliquots of the concentrated stocks were subsequently diluted in a two-step process: first in DMSO and then in protein equilibration buffer (10 mM HEPES, 100 mM NaCl, pH 7.5, 0.01%/0.001% LMNG/CHS), yielding ligand solutions at a final concentration of 1.122 mM and 43% DMSO. GLP-1 peptides were prepared using a procedure analogous to that employed for the compounds, with the following modifications. Peptides were dissolved in DMSO to generate initial stock solutions at a concentration of 5 mM. Aliquots of each peptide stock were further diluted in DMSO to a concentration of 750 µM and subsequently mixed at a 1:1.5 ratio with protein equilibration buffer containing 0.167% bacitracin, yielding ligand solutions at a final concentration of 300 µM and 40% DMSO.

Mixing 1 µL of 1.122 mM compound solutions (danuglipron and Chu-128) with 7.5 µL of GLP-1R-Gs (15 µM) produced a 10-fold molar excess of ligand and a final DMSO concentration of 5%. Similarly, mixing 1 µL of 300 µM GLP-1 peptide solutions (GLP-1[7-36] and GLP-1[9-36],) with the GLP-1R–Gs sample (16 µM) resulted in a 2.5-fold molar excess of ligand and final DMSO and bacitracin concentrations of 5% and 0.02%, respectively. In both cases, the mixing procedure ensured saturating ligand conditions for following HDX-MS experiments. Both DMSO and bacitracin were also included in the apo GLP-1R–Gs samples to ensure consistency with the conditions used for the ligand-bound samples.

To ensure sufficient ligand-binding saturation under deuterium exchange conditions, the GLP-1R–Gs complex was incubated with a 10-fold molar excess of ligand for 1 h (45 min at 7 °C followed by 15 min at room temperature). This incubation allowed both ligand binding and temperature equilibration prior to deuterium exchange experiments.

### Hydrogen/deuterium mass spectrometry

Measurements were performed on an ACQUITY UPLC M-Class System with HDX Technology (Waters) directly coupled to a Xevo G2-XS QToF Mass Spectrometer (Waters). with manual injection of all samples and trapping and separation system maintained at 0 °C. HDX labelling was performed as follows. In danuglipron and Chu-128 experiments, 8.5 µL of GLP-1R–Gs sample, either in the apo state or pre-incubated with ligand, was diluted with 35.75 µL of labelling buffer (10 mM HEPES, 100 mM NaCl, pD 7.1, 0.01%/0.001% LMNG/CHS) at 20 °C, resulting in 81% D₂O labelling. Samples were incubated for 10, 50 (NPAs only), 1250, and 6250 s and subsequently quenched by the addition of quench buffer (100 mM NaH₂PO₄, 7 M urea, 585 mM TCEP, pH 2.3) in a volume equivalent to that of the labelling buffer. Quenched samples were immediately frozen in liquid nitrogen without any pre-incubation prior to freezing. In the quenched samples, urea and TCEP were present at final concentrations of 3 M and 256 mM, respectively, with no other detergents added to aid protein unfolding. Frozen samples were thawed prior to manual injection and 80 µL of the quenched sample was then loaded onto a 50 µL sample loop before being injected onto an online system of PNGase deglycosylation column followed by Nepenthesin-II digestion column (PNGase-Nepenthesin II system, (Affipro)) maintained at room temperature and passed through by solvent A (0.23% formic acid in water) at flow rate of 200 µL/min. Generated peptides were trapped onto Acquity UPLC BEH C18 VanGuard pre-column (1.7 µm, 2.1 mm x 5 mm (Waters)) for 3 min, and then eluted and separated on Acquity UPLC BEH C18 analytical column (1.7 µm, 1.0 mm x 100 mm (Waters)) with a linear gradient over 7.5 minutes from 8 to 35% solvent B (0.23% formic acid in acetonitrile) at a flow rate of 40 µL/min and at 0 °C. To prevent peptide carryover, PNGase-Nepenthesin II system was washed twice during the linear gradient using a protease wash solution (1.6 M guanidine-HCl, 4% acetonitrile, 0.8% formic acid and 0.1% n-Dodecylphosphocholine (Fos-Choline-12), pH 2.5). Further, the chromatographic columns were washed after each sample run using a sawtooth gradient, with an additional washing step applied to PNGase-Nepenthesin II system. The eluted peptides were ionized by electrospray into the Xevo G2-XS QToF mass spectrometer. MS^E^ data was acquired in positive ion mode with a ramped collision energy from 20 to 45 V. Sodium iodide and leucine enkephalin were used for calibration and mass accuracy correction, respectively.

All deuterium time points and controls were performed in triplicate. Sequence identification was performed from MS^E^ data of digested undeuterated samples of GLP-1R-Gs using the ProteinLynx Global Server 2.5.1 software. The output peptides were then filtered using DynamX (v. 3.0) using the following filtering parameters: minimum intensity of 3000, maximum peptide sequence length of 30, minimum products per amino acid of 0.1, minimum score of 8, maximum MH^+^ error threshold of 10 ppm, file threshold of 2 out of 3 and retention time RSD of 5%. Additionally, all spectra were visually inspected and only those with a suitable signal to noise ratios were used for analysis. The amount of relative deuterium uptake for each peptide was determined using DynamX (v. 3.0) and was not corrected for back exchange (83).

Confidence intervals for differential HDX-MS (ΔHDX) measurements of any individual time point, as well as the aggregated differences (∑ΔHDX) over all timepoints, were determined according to Houde et al. (84) using Deuteros software (v. 1.0; 49, 84). Significant peptides were considered only those which satisfied a ΔHDX confidence interval of 98%. All ΔHDX GLP-1R-Gs structure figures were generated from the data using PyMOL Molecular Graphics System (v. 3.1.6.1.; 85) utilising AlphaFold 3 predicted structure or PDB structures of GLP-1R 6X19.

## Data availability

The HDX-MS data generated in this study have been provided in the Source data file, and the HDX-MS summary tables and uptake plots have been provided as Supplementary Data. Furthermore, the mass spectrometry proteomics data have been deposited to the MassIVE partner repository via the ProteomeXchange Consortium with the dataset identifier PXD081918.

## Acknowledgements

The authors thank Kris Conners, Marc Rutter, Russell Madsen, Brad Condon, Peggy Kearins, Marijane Russell, and Kevin Bain for their contributions to molecular biology, protein expression, and biochemistry. The authors thank UKRI and Eli Lilly for funding. JS and ER were supported by Eli Lilly and Company through the Lilly Research Award Program. ER is supported by a UKRI Future Leaders Fellowship (MR/S015426/1 and MR/X009580/1).

## Author contributions

**Jakub Sýs:** Methodology, Validation, Formal Analysis, Investigation (all in Purification of GLP-1R:Gs non-covalent complex, HDX-MS experiments), Data Curation (HDX-MS experiments), Writing - Original Draft, Visualization

**Joseph D Ho:** Conceptualization, Methodology (Purification of the apo-active GLP-1R-Gαs-Gβγ complex), Resources, Supervision

**Aaron D Showalter:** Methodology, Investigation, Validation, Visualization (all in Cyclic AMP accumulation assays)

**An-Ping Yu:** Methodology, Investigation, Validation, Visualization (all in Cyclic AMP accumulation assays)

**David B Wainscott:** Methodology, Investigation, Validation, Visualization (all in Radioligand binding)

**Veronica Laos:** Methodology, Investigation, Validation, Visualization (all in Affinity Selection Mass Spectrometry (ASMS) assay)

**Howard Broughton:** Conceptualization, Supervision

**Kyle W Sloop:** Conceptualization, Writing - Review & Editing, Supervision, Project administration

**Alfonso Espada:** Conceptualization, Writing - Review & Editing, Supervision, Project administration, Funding acquisition

**Eamonn Reading:** Conceptualization, Resources, Writing - Review & Editing, Supervision, Funding acquisition

## References

1. B. Ahrén, Islet G protein-coupled receptors as potential targets for treatment of type 2 diabetes. Nat. Rev. Drug Discov. 8, 369–385 (2009).

2. L. F. Kolakowski, GCRDb: A G-Protein–Coupled Receptor. Receptors Channels 2, 1–7 (1994).

3. R. Fredriksson, M. C. Lagerström, L. G. Lundin, H. B. Schiöth, The G-protein-coupled receptors in the human genome form five main families. Phylogenetic analysis, paralogon groups, and fingerprints. Mol. Pharmacol. 63, 1256–1272 (2003).

4. Y. M. Cho, C. E. Merchant, T. J. Kieffer, Targeting the glucagon receptor family for diabetes and obesity therapy. Pharmacol. Ther. 135, 247–278 (2012).

5. P. L. Brubaker, D. J. Drucker, Structure-Function of the Glucagon Receptor Family of G Protein-Coupled Receptors: The Glucagon, GIP, GLP-1, and GLP-2 Receptors. Receptors Channels 8, 179–188 (2002).

6. C. d. Graaf et al., Glucagon-Like Peptide-1 and Its Class B G Protein–Coupled Receptors: A Long March to Therapeutic Successes. Pharmacol. Rev. 68, 954–1013 (2016).

7. J. Gromada, J. J. Holst, P. Rorsman, Cellular regulation of islet hormone secretion by the incretin hormone glucagon-like peptide 1. Pflügers Archiv 435, 583–594 (1998).

8. E. Renström, L. Eliasson, P. Rorsman, Protein kinase A-dependent and-independent stimulation of exocytosis by cAMP in mouse pancreatic B-cells. The Journal of physiology 502, 105 (1997).

9. B. Kreymann, M. A. Ghatei, G. Williams, S. R. Bloom, Glucagon-like peptide-1 7-36: a physiological incretin in man. The Lancet 330, 1300–1304 (1987).

10. S. Mojsov, G. C. Weir, J. F. Habener, Insulinotropin: glucagon-like peptide I (7-37) co-encoded in the glucagon gene is a potent stimulator of insulin release in the perfused rat pancreas. The Journal of Clinical Investigation 79, 616–619 (1987).

11. J. Gromada, B. Brock, O. Schmitz, P. Rorsman, Glucagon-like peptide-1: regulation of insulin secretion and therapeutic potential. Basic Clin. Pharmacol. Toxicol. 95, 252–262 (2004).

12. A. Wettergren et al., Truncated GLP-1 (proglucagon 78–107-amide) inhibits gastric and pancreatic functions in man. Dig. Dis. Sci. 38, 665–673 (1993).

13. L. L. Baggio, Q. Huang, T. J. Brown, D. J. Drucker, A Recombinant Human Glucagon-Like Peptide (GLP)-1–Albumin Protein (Albugon) Mimics Peptidergic Activation of GLP-1 Receptor–Dependent Pathways Coupled With Satiety, Gastrointestinal Motility, and Glucose Homeostasis. Diabetes 53, 2492–2500 (2004).

14. L. L. Baggio, D. J. Drucker, Biology of incretins: GLP-1 and GIP. Gastroenterology 132, 2131–2157 (2007).

15. S. E. Kanoski, S. M. Fortin, M. Arnold, H. J. Grill, M. R. Hayes, Peripheral and central GLP-1 receptor populations mediate the anorectic effects of peripherally administered GLP-1 receptor agonists, liraglutide and exendin-4. Endocrinology 152, 3103–3112 (2011).

16. R. Mentlein, B. Gallwitz, W. E. Schmidt, Dipeptidyl-peptidase IV hydrolyses gastric inhibitory polypeptide, glucagon-like peptide-1(7-36)amide, peptide histidine methionine and is responsible for their degradation in human serum. Eur. J. Biochem. 214, 829–835 (1993).

17. C. F. Deacon, A. H. Johnsen, J. J. Holst, Degradation of glucagon-like peptide-1 by human plasma in vitro yields an N-terminally truncated peptide that is a major endogenous metabolite in vivo. J. Clin. Endocrinol. Metab. 80, 952–957 (1995).

18. S. Pearson, N. Kietsiriroje, R. A. Ajjan, Oral semaglutide in the management of type 2 diabetes: a report on the evidence to date. Diabetes, Metabolic Syndrome and Obesity, 2515-2529 (2019).

19. G. Singh, M. Krauthamer, M. Bjalme-Evans, Wegovy (semaglutide): a new weight loss drug for chronic weight management. J Investig Med 70, 5–13 (2022).

20. T. J. Moretto et al., Efficacy and tolerability of exenatide monotherapy over 24 weeks in antidiabetic drug—naive patients with type 2 diabetes: A randomized, double-blind, placebo-controlled, parallel-group study. Clinical therapeutics 30, 1448–1460 (2008).

21. A. Astrup et al., Effects of liraglutide in the treatment of obesity: a randomised, double-blind, placebo-controlled study. The Lancet 374, 1606–1616 (2009).

22. I. The Medical Letter, Tirzepatide (Mounjaro) for type 2 diabetes. Medical Letter on Drugs and Therapeutics 64, 105–107 (2022).

23. L. Collins, R. A. Costello, "Glucagon-Like Peptide-1 Receptor Agonists" in StatPearls. (StatPearls Publishing Copyright © 2026, StatPearls Publishing LLC., Treasure Island (FL) ineligible companies., 2026).

24. I. Pfizer, Pfizer provides update on oral GLP-1 receptor agonist danuglipron. (2025).

25. S. Madsbad, Review of head-to-head comparisons of glucagon-like peptide-1 receptor agonists. Diabetes Obes Metab 18, 317–332 (2016).

26. L. Chugai Pharmaceutical Co (2018) Pyrazolopyridine derivative having GLP-1 receptor agonist effect.

27. S. Wharton et al., Orforglipron, an Oral Small-Molecule GLP-1 Receptor Agonist for Obesity Treatment. *New Engl*. J. Med. 393, 1796–1806 (2025).

28. X. Zhang et al., Differential GLP-1R Binding and Activation by Peptide and Non-peptide Agonists. Mol. Cell 80, 485–500.e487 (2020).

29. T. Kawai et al., Structural basis for GLP-1 receptor activation by LY3502970, an orally active nonpeptide agonist. Proc Natl Acad Sci U S A 117, 29959–29967 (2020).

30. A. S. Hauser, M. M. Attwood, M. Rask-Andersen, H. B. Schiöth, D. E. Gloriam, Trends in GPCR drug discovery: new agents, targets and indications. Nat. Rev. Drug Discov. 16, 829–842 (2017).

31. R. Santos et al., A comprehensive map of molecular drug targets. Nat. Rev. Drug Discov. 16, 19–34 (2017).

32. K. Sriram, P. A. Insel, G Protein-Coupled Receptors as Targets for Approved Drugs: How Many Targets and How Many Drugs? Mol. Pharmacol. 93, 251–258 (2018).

33. P. Conflitti et al., Functional dynamics of G protein-coupled receptors reveal new routes for drug discovery. Nature Reviews Drug Discovery 24, 251–275 (2025).

34. C. J. Fairweather et al., Conformational Dynamics of Amylin Receptors Revealed by Hydrogen–Deuterium Exchange Mass Spectrometry. J. Am. Chem. Soc. 148, 16702–16712 (2026).

35. H. E. Kato et al., Conformational transitions of a neurotensin receptor 1–Gi1 complex. Nature 572, 80–85 (2019).

36. A. A. Vo et al., Snapshots of the dynamic basis of NTSR1 G protein subtype promiscuity. Nature 652, 803–811 (2026).

37. E. I. James, T. A. Murphree, C. Vorauer, J. R. Engen, M. Guttman, Advances in Hydrogen/Deuterium Exchange Mass Spectrometry and the Pursuit of Challenging Biological Systems. Chemical Reviews 122, 7562–7623 (2022).

38. G. R. Masson et al., Recommendations for performing, interpreting and reporting hydrogen deuterium exchange mass spectrometry (HDX-MS) experiments. Nat. Methods 16, 595–602 (2019).

39. V. Vinciauskaite, G. R. Masson, Fundamentals of HDX-MS. Essays Biochem 67, 301–314 (2023).

40. D. M. Ferraro, N. D. Lazo, A. D. Robertson, EX1 Hydrogen Exchange and Protein Folding. Biochemistry 43, 587–594 (2004).

41. D. D. Weis, T. E. Wales, J. R. Engen, M. Hotchko, L. F. Ten Eyck, Identification and characterization of EX1 kinetics in H/D exchange mass spectrometry by peak width analysis. J. Am. Soc. Mass Spectrom. 17, 1498–1509 (2006).

42. N. M. Duc et al., Effective Application of Bicelles for Conformational Analysis of G Protein-Coupled Receptors by Hydrogen/Deuterium Exchange Mass Spectrometry. J. Am. Soc. Mass Spectrom. 26, 808–817 (2015).

43. K. Krishna Kumar et al., Negative allosteric modulation of the glucagon receptor by RAMP2. Cell 186, 1465–1477.e1418 (2023).

44. Y. Zhang et al., Cryo-EM structure of the activated GLP-1 receptor in complex with a G protein. Nature 546, 248–253 (2017).

45. M. Casiraghi et al., Structure and dynamics determine G protein coupling specificity at a class A GPCR. Science Advances 11, eadq3971 (2025).

46. I. R. Möller et al., Probing the conformational impact of detergents on the integral membrane protein LeuT by global HDX-MS. Journal of Proteomics 225, 103845 (2020).

47. P. Zhao et al., Activation of the GLP-1 receptor by a non-peptidic agonist. Nature 577, 432–436 (2020).

48. E. Wolf et al., Quantitative Hydrogen–Deuterium Exchange Mass Spectrometry for Simultaneous Structural Characterization and Affinity Indexing of Single Target Drug Candidate Libraries. Analytical Chemistry 96, 13015–13024 (2024).

49. A. M. C. Lau, Z. Ahdash, C. Martens, A. Politis, Deuteros: software for rapid analysis and visualization of data from differential hydrogen deuterium exchange-mass spectrometry. Bioinformatics 35, 3171–3173 (2019).

50. J. Lesne et al., Conformational maps of human 20S proteasomes reveal PA28- and immuno-dependent inter-ring crosstalks. Nat. Commun. 11, 6140 (2020).

51. D. G. Lambright, J. P. Noel, H. E. Hamm, P. B. Sigler, Structural determinants for activation of the α-subunit of a heterotrimeric G protein. Nature 369, 621–628 (1994).

52. S. R. Sprang, Invited review: Activation of G proteins by GTP and the mechanism of Gα-catalyzed GTP hydrolysis. Biopolymers 105, 449–462 (2016).

53. G. T. Hammons, R. M. Smith, L. Jarett, Inhibition by bacitracin of rat adipocyte plasma membrane degradation of 125I-insulin is associated with an increase in plasma membrane bound insulin and a potentiation of glucose oxidation by adipocytes. J. Biol. Chem. 257, 11563–11570 (1982).

54. Q. Xiao, W. Jeng, M. B. Wheeler, Characterization of glucagon-like peptide-1 receptor-binding determinants. Journal of Molecular Endocrinology 25, 321–335 (2000).

55. A. K. Nielsen et al., Substrate-induced conformational dynamics of the dopamine transporter. Nat. Commun. 10, 2714 (2019).

56. P. S. Merkle et al., Substrate-modulated unwinding of transmembrane helices in the NSS transporter LeuT. Science Advances 4, eaar6179.

57. A. S. Hauser et al., GPCR activation mechanisms across classes and macro/microscales. Nat. Struct. Mol. Biol. 28, 879–888 (2021).

58. Y. H. Feng, S. S. Karnik, Role of transmembrane helix IV in G-protein specificity of the angiotensin II type 1 receptor. J. Biol. Chem. 274, 35546–35552 (1999).

59. F. Wu et al., Full-length human GLP-1 receptor structure without orthosteric ligands. Nat Commun 11, 1272 (2020).

60. H. N. Do, A. Haldane, R. M. Levy, Y. Miao, Unique features of different classes of G-protein-coupled receptors revealed from sequence coevolutionary and structural analysis. Proteins 90, 601–614 (2022).

61. A. Mafi, S. K. Kim, W. A. Goddard, 3rd, The mechanism for ligand activation of the GPCR-G protein complex. Proc Natl Acad Sci U S A 119, e2110085119 (2022).

62. D. Hilger et al., Structural insights into differences in G protein activation by family A and family B GPCRs. Science 369 (2020).

63. Y. L. Liang et al., Toward a Structural Understanding of Class B GPCR Peptide Binding and Activation. Mol. Cell 77, 656–668.e655 (2020).

64. G. Mattedi, S. Acosta-Gutiérrez, T. Clark, F. L. Gervasio, A combined activation mechanism for the glucagon receptor. Proc Natl Acad Sci U S A 117, 15414–15422 (2020).

65. Z. Lawson, M. Wheatley, The third extracellular loop of G-protein-coupled receptors: more than just a linker between two important transmembrane helices. Biochem. Soc. Trans. 32, 1048–1050 (2004).

66. Z. L. Lu, J. W. Saldanha, E. C. Hulme, Seven-transmembrane receptors: crystals clarify. Trends Pharmacol. Sci. 23, 140–146 (2002).

67. F. M. Décaillot et al., Opioid receptor random mutagenesis reveals a mechanism for G protein-coupled receptor activation. Nat. Struct. Biol. 10, 629–636 (2003).

68. M. M. Zhao, R. J. Gaivin, D. M. Perez, The third extracellular loop of the beta2- adrenergic receptor can modulate receptor/G protein affinity. Mol. Pharmacol. 53, 524–529 (1998).

69. S. Kosugi, T. Mori, The third exoplasmic loop of the thyrotropin receptor is partially involved in signal transduction. FEBS Lett. 349, 89–92 (1994).

70. K. Palczewski et al., Crystal structure of rhodopsin: A G protein-coupled receptor. Science 289, 739–745 (2000).

71. K. Ohyama et al., Disulfide bridges in extracellular domains of angiotensin II receptor type IA. Regul Pept 57, 141–147 (1995).

72. Y.-L. Liang et al., Phase-plate cryo-EM structure of a class B GPCR–G-protein complex. Nature 546, 118–123 (2017).

73. Y. L. Liang et al., Phase-plate cryo-EM structure of a biased agonist-bound human GLP- 1 receptor-Gs complex. Nature 555, 121–125 (2018).

74. G. M. West et al., Glucagon-Like Peptide-1 Receptor Ligand Interactions: Structural Cross Talk between Ligands and the Extracellular Domain. PLOS ONE 9, e105683 (2014).

75. P. M. Scrosati, V. Yin, L. Konermann, Hydrogen/Deuterium Exchange Measurements May Provide an Incomplete View of Protein Dynamics: a Case Study on Cytochrome c. Analytical Chemistry 93, 14121–14129 (2021).

76. S. Maeda et al., Development of an antibody fragment that stabilizes GPCR/G-protein complexes. Nat Commun 9, 3712 (2018).

77. S. G. Rasmussen et al., Crystal structure of the β2 adrenergic receptor-Gs protein complex. Nature 477, 549–555 (2011).

78. H.-Y. Yen et al., PtdIns(4,5)P2 stabilizes active states of GPCRs and enhances selectivity of G-protein coupling. Nature 559, 423–427 (2018).

79. K. W. Chao et al., Human class B1 GPCR modulation by plasma membrane lipids. Communications Biology 9, 317 (2026).

80. R. Baccouch, E. Rascol, K. Stoklosa, I. D. Alves, The role of the lipid environment in the activity of G protein coupled receptors. Biophys. Chem. 285, 106794 (2022).

81. C. Guffick, A. Politis, HDX-MS in micelles and membranes for small molecule and biopharmaceutical development. Curr. Opin. Struct. Biol. 94, 103077 (2025).

82. F. S. Willard, et al., Tirzepatide is an imbalanced and biased dual GIP and GLP-1 receptor agonist. JCI Insight 5 (2020).

83. T. E. Wales, M. J. Eggertson, J. R. Engen, "Considerations in the Analysis of Hydrogen Exchange Mass Spectrometry Data" in Mass Spectrometry Data Analysis in Proteomics, R. Matthiesen, Ed. (Humana Press, Totowa, NJ, 2013), 10.1007/978-1-62703-392-3_11, pp. 263–288.

84. D. Houde, S. A. Berkowitz, J. R. Engen, The Utility of Hydrogen/Deuterium Exchange Mass Spectrometry in Biopharmaceutical Comparability Studies. J. Pharm. Sci. 100, 2071–2086 (2011).

85. L. L. C. Schrödinger (2010) The PyMOL molecular graphics system.

