## Supplementary Information for "Structural dynamics underlying agonist activation of a GLP-1R-Gs precoupled complex"

### SI appendix

**Supplementary Discussion**

*Initial attempts to purify noncovalent, ligand-free state of GLP-1R:$G_s$*

Here, we aimed to elucidate the dynamics of GLP-1R in complex with $G_s$ and to determine agonist-induced structural perturbations by HDX-MS. HDX-MS is best achieved by performing differential experiments (ΔHDX) that compare the protein in both its ligand-free (apo) and ligand-bound (complex) states. When purified in the absence of ligand, GLP-1R solubilized in LMNG/CHS and immobilized through its N-terminal FLAG tag exhibited low purity and evidence of partial truncation (**Fig. S1A-D**). We then established an alternative purification protocol utilising a 6xHis tag at Gγ C-terminus for cobalt-based IMAC (Talon resin) followed by two-step size exclusion chromatography (SEC). Using this protocol, we obtained preliminary HDX-MS data for the non-covalent GLP-1R:Gs complex in its ligand-free state identifying 47 peptides, yielding 75.2% sequence coverage of GLP-1R with an average redundancy of only 1.46, with the deuteration profiles indicating presence of properly folded GLP-1R (**Fig. S1E**). Despite the initial success, the persistent instability of the construct in the ligand-free state led to low peptide yield and intensity, preventing a thorough HDX-MS study using noncovalent apo GLP-1R:$G_s$.

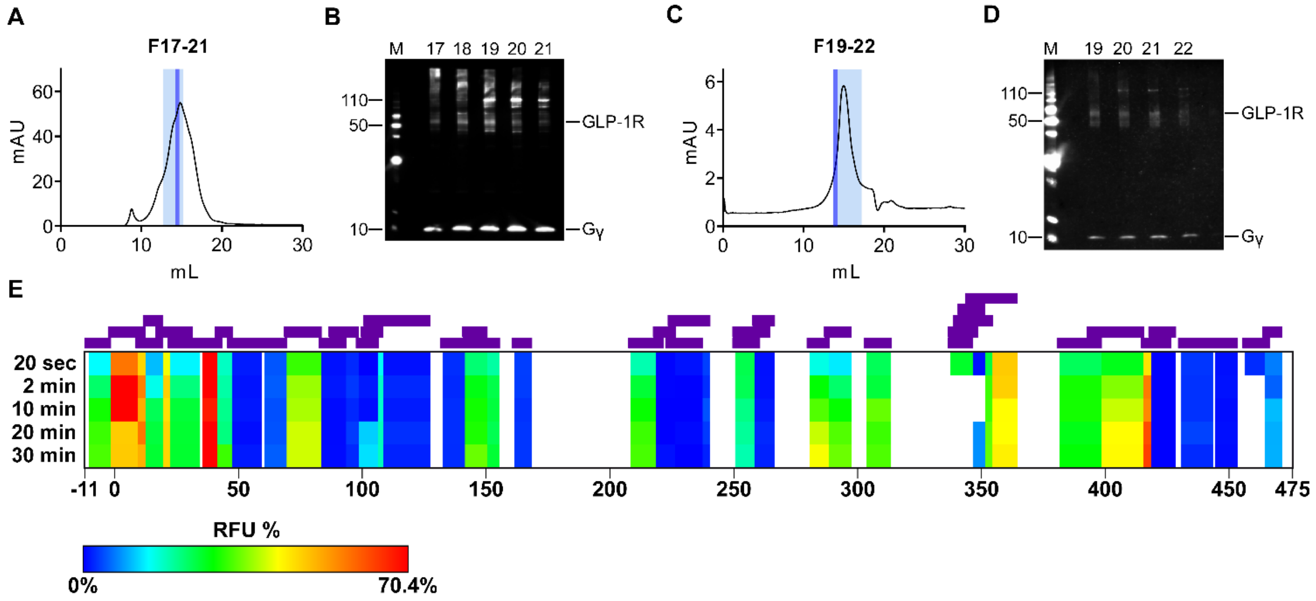


**Figure S1. Purification procedure and Conformational Dynamics of GLP-1R in a Noncovalent Complex with Gs Protein.** (A) Size-exclusion chromatography of the GLP-1R:Gs sample eluted with 150 mM imidazole; fractions 17–21 are shown in light blue, with fraction 20 highlighted in dark blue. (B) Immunoblot of fractions 17–21 using a monoclonal anti-6His antibody, showing Gsγ (6.5 kDa) and GLP-1R (50–60 kDa). A PageRuler His-tagged protein ladder (Thermo) was used as a marker. (C) Size-exclusion chromatography of fraction 20, with fractions 19–22 shown in light blue and fraction 19 highlighted in dark blue. (D) Immunoblot of fractions 19–22 using a monoclonal anti-6His antibody, showing Gγ (6.5 kDa) and GLP-1R (50–60 kDa). Fraction 19 was used for HDX-MS. (E) HDX MS fractional uptake values for apo GLP-1R in a noncovalent complex with the Gs protein over deuteration time points ranging from 20 seconds to 30 minutes. Peptide-level resolution is shown, with the positions of the 43 identified peptides indicated by purple bars above the heatmap, corresponding to 75.2% sequence coverage and an average redundancy of 1.46. HDX MS data were collected in a single replicate. To ensure consistency with UniProt protein sequence numbering, residue numbering starts at −11; thus, the signal peptide and FLAG tag sequences are assigned negative residue numbers. The raw data, along with the DynamX file, have been deposited in the PRIDE repository.

(a)

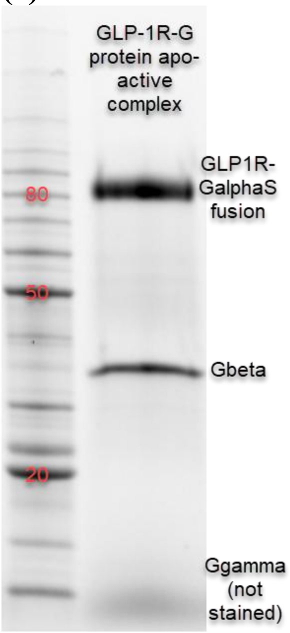


(b)

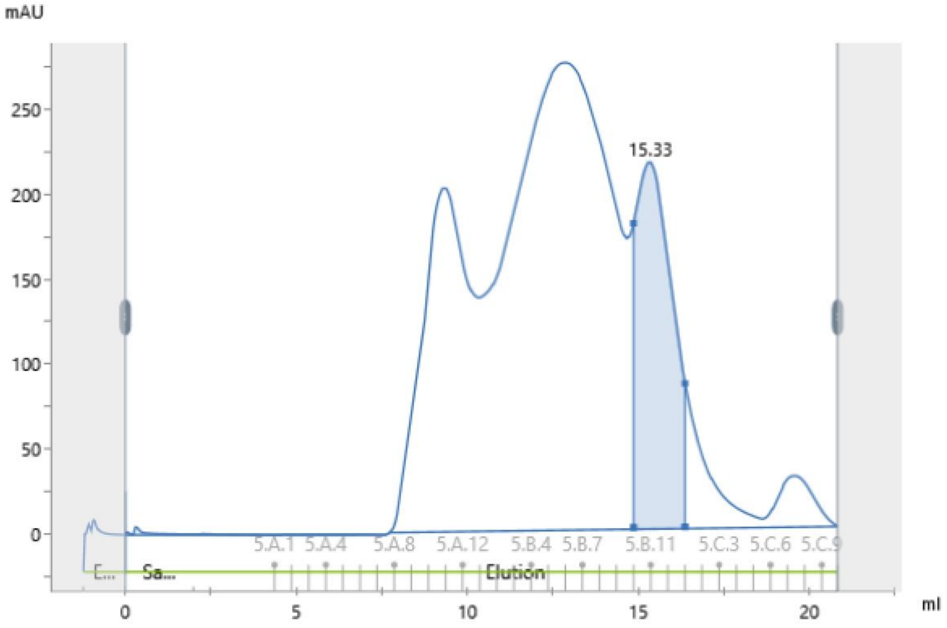


**Figure S2. The protein biochemistry of GLP-1R–Gs + Gbeta/gamma complex. (a) SDS-PAGE in BioRad tryptophan fluorescence gel and (b) Superose 6 size-exclusion chromatography.**

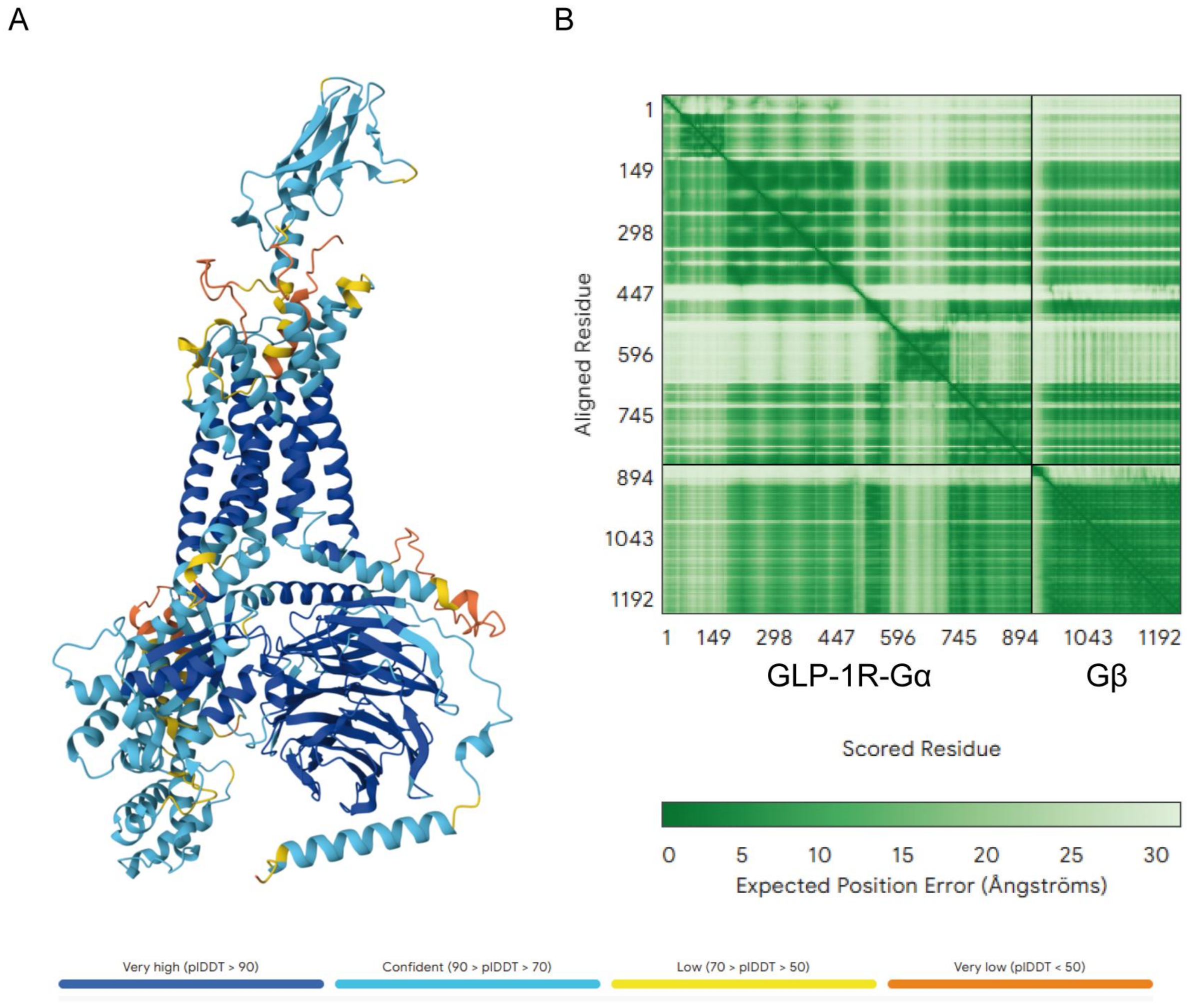


**Figure S3. AlphaFold prediction of GLP-1R-Gs (GLP-1R-Gα:Gβ) structure. (A)** Predicted structure of GLP-1R-Gs, with GLP-1R-Gα (chain A) and Gβ (chain B). Coloured coded based on the per-residue confidence score (pLDDT). Residues with very high scores (pLDDT > 90), confident scores (pLDDT > 70), low scores (pLDDT > 50), and very low scores (pLDDT < 50) are indicated in dark blue, light blue, yellow, and orange, respectively. A low confidence score (<50) is likely to indicate unstructured regions. **(B)** Predicted aligned error plot. The AlphaFold's expected position error at residue X is indicated by colour position at (x, y), when the predicted and true structures are aligned on residue y. The colour bar indicates the confidence level of AlphaFold's prediction, with dark green representing high confidence and light green indicating low confidence. A black line within the error plot delimits the chain borders.

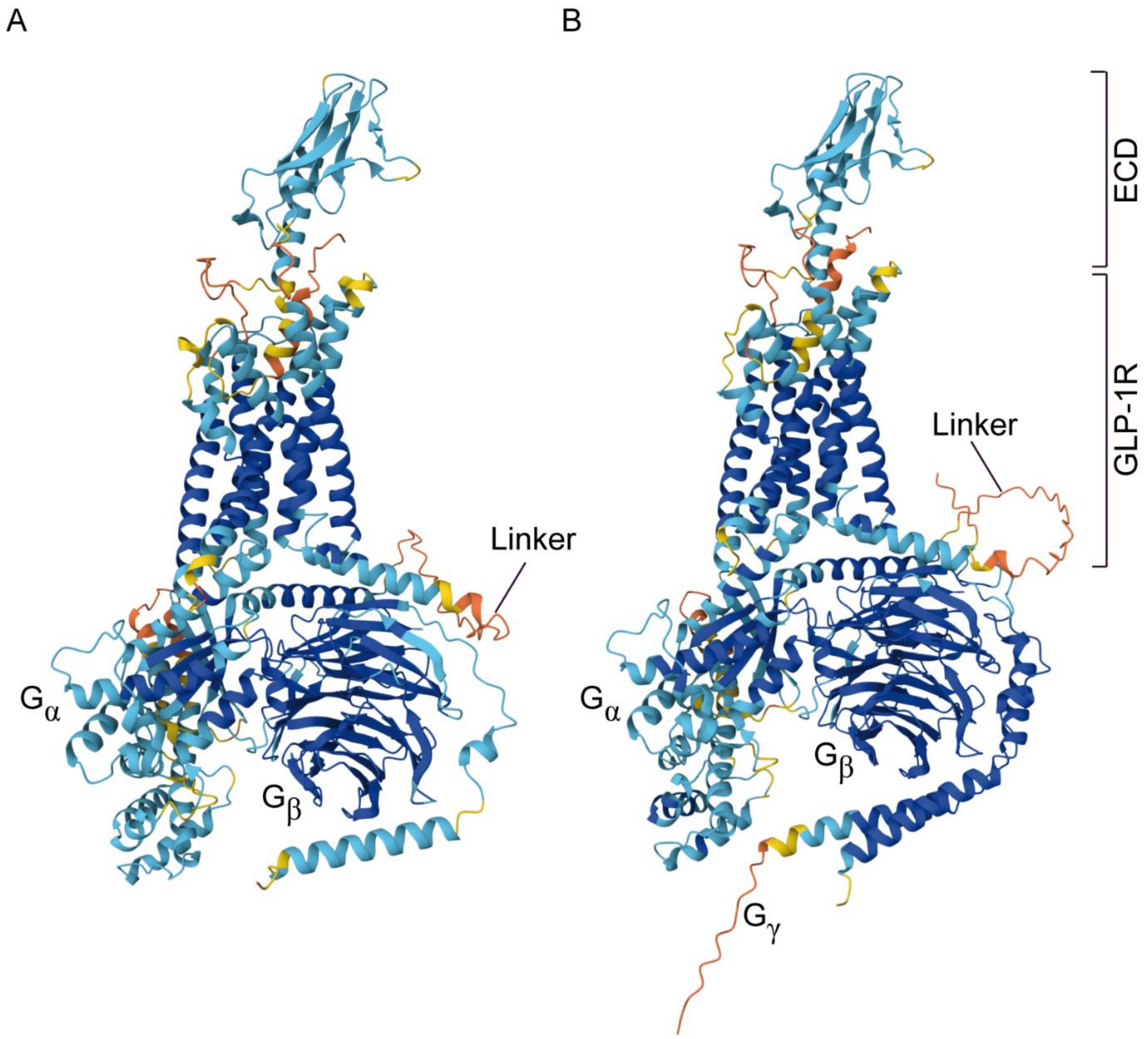


**Figure S4. Comparison of AlphaFold predicted structures of GLP-1R-Gα:Gβ (GLP-1R-Gs) and GLP-1R-Gα:Gβ:Gγ. (A)** Predicted structure of GLP-1R-Gs, with GLP-1R-Gα (chain A) and Gβ (chain B). **(B)** Predicted structure of GLP-1R-Gs, with GLP-1R-Gα (chain A), Gβ (chain B) and Gγ (chain C). Structures are colour-coded based on the per-residue confidence score (pLDDT). Residues with very high scores (pLDDT > 90), confident scores (pLDDT > 70), low scores (pLDDT > 50), and very low scores (pLDDT < 50) are indicated in dark blue, light blue, yellow, and orange, respectively. Neither GLP-1R, Gα, nor Gβ exhibit differences in their predicted conformations with respect to the absence or presence of the Gγ subunit of the Gs protein. The only exception is the linker region bridging GLP-1R and Gα in the GLP-1R-Gs covalent construct, which is indicated as a region with a low confidence score in both structures. For simplicity, and because no HDX-MS data were collected for Gγ, the predicted GLP-1R-Gα:Gβ complex, hereafter referred to as GLP-1R–Gs, was used to visualise HDX-MS data where appropriate.

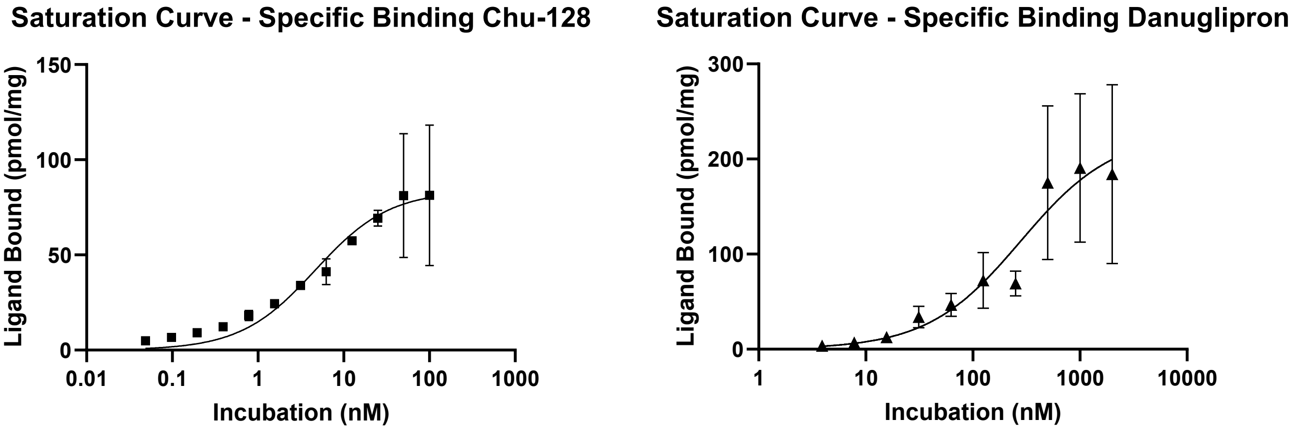


**Figure S5. High-affinity equilibrium dissociation constants ($K_D$) for Chu-128 and Danuglipron determined using an Affinity Selection Mass Spectrometry (ASMS) assay.** Equilibrium dissociation constants ($K_D$) of 4.63 nM and 279.5 nM were obtained for Chu-128 (LSN3955529) and Danuglipron (LSN3535219), respectively, based on triplicate measurements. GLP-1R–Gs was incubated with a range of concentrations of Chu-128 (left) and Danuglipron (right). The plots illustrate the bound ligand quantified versus the incubation concentration.

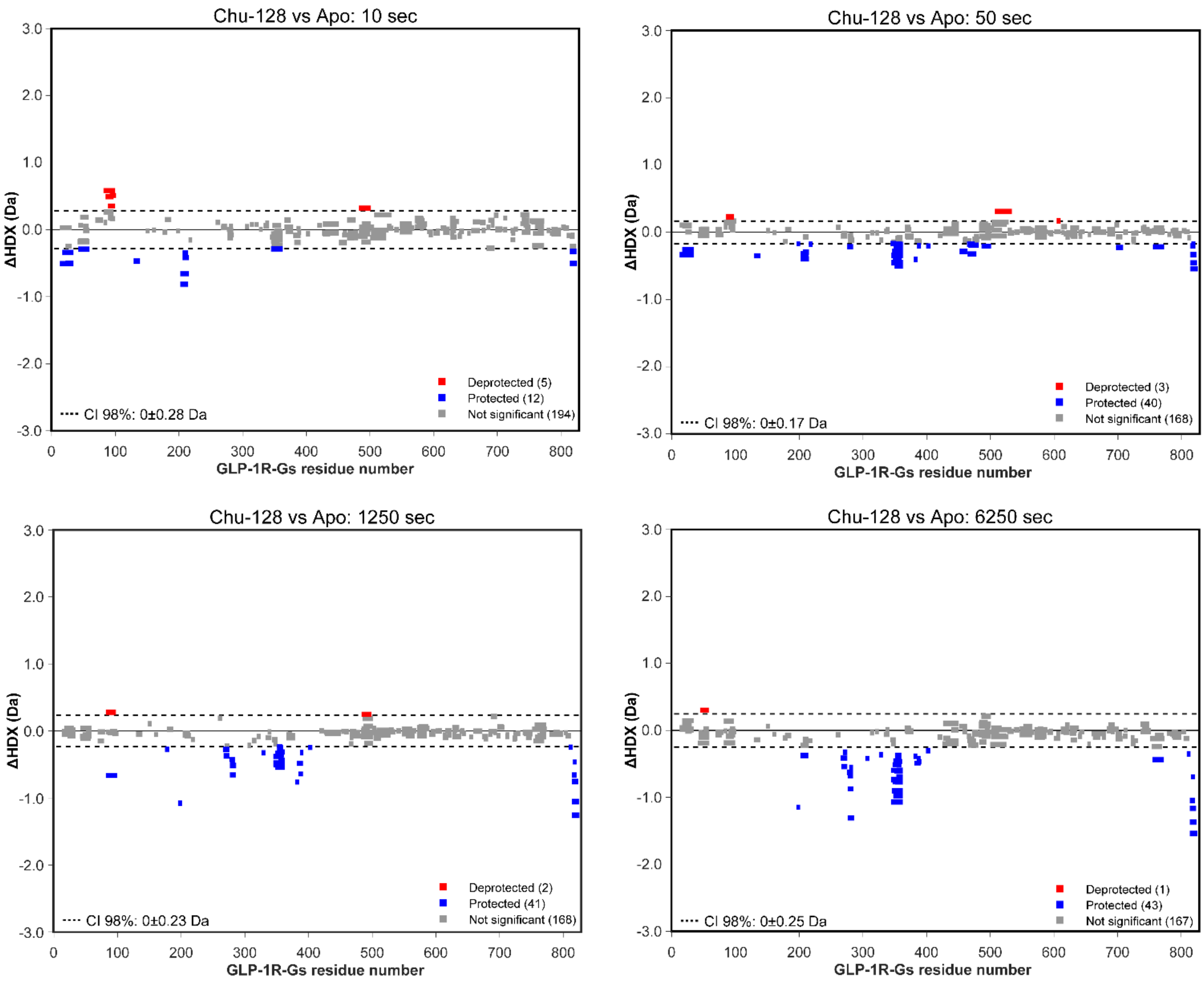


**Figure S6. Individual Woods plots for two-state comparison of Chu-128 vs Apo.** Differential HDX (ΔHDX) plots from a two-state comparison of the Chu-128-bound state versus the apo state at individual time points (10, 50, 1250, and 6250 seconds). Red signifies peptides with increased HDX between states, while blue represents peptides with decreased HDX. The 98% confidence intervals are shown as black dashed lines, and grey data denote peptides with insignificant ΔHDX. All measurements were performed in triplicate.

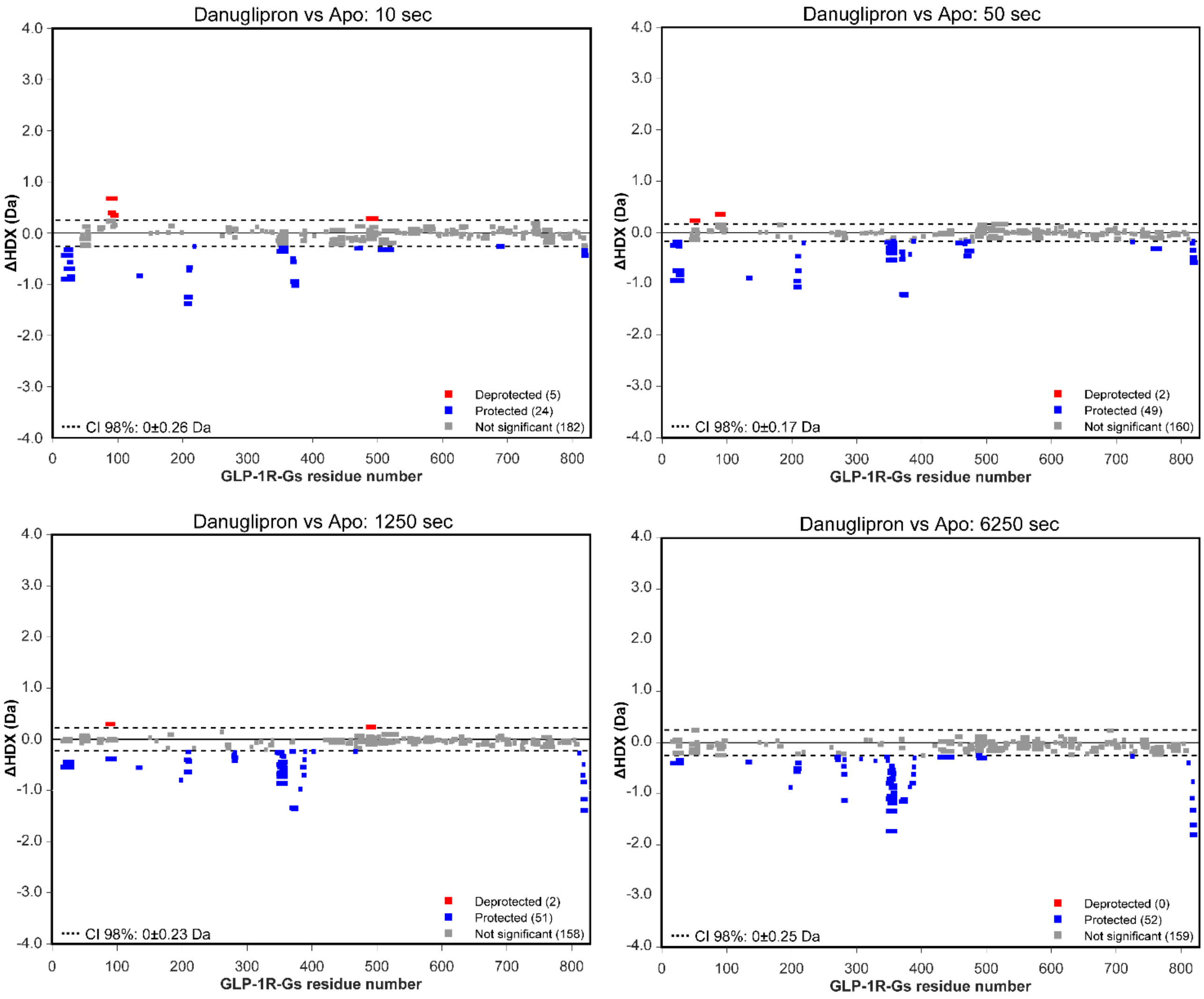


**Figure S7. Individual Woods plots for two-state comparison of Danu vs Apo.** Differential HDX (ΔHDX) plots from a two-state comparison of the Danuglipron-bound state versus the apo state at individual time points (10, 50, 1250, and 6250 seconds). Red signifies peptides with increased HDX between states, while blue represents peptides with decreased HDX. The 98% confidence intervals are shown as black dashed lines, and grey data denote peptides with insignificant ΔHDX. All measurements were performed in triplicate.

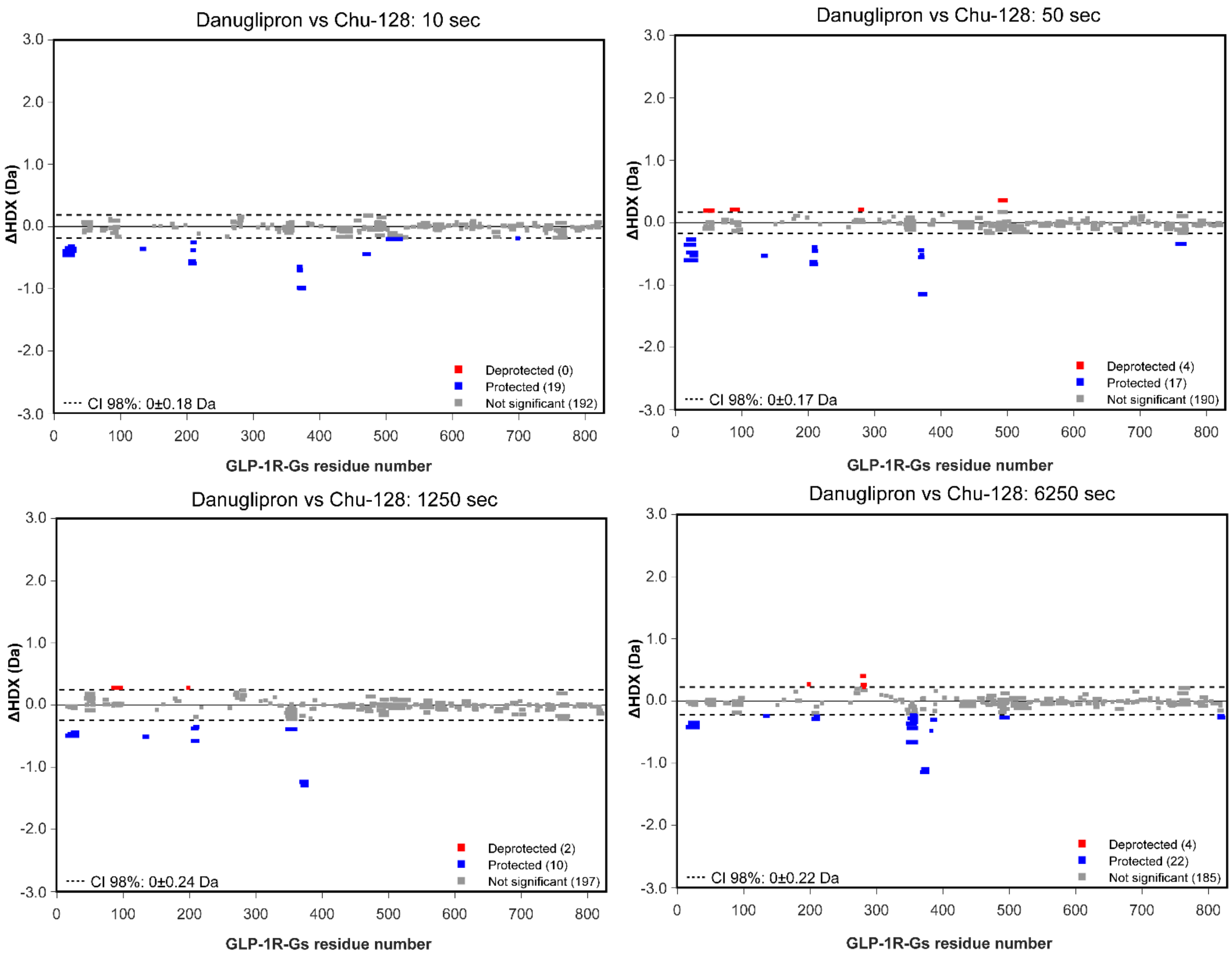


**Figure S8. Individual Woods plots for two-state comparison of Danuglipron vs Chu-128.** Differential HDX (ΔHDX) plots from a two-state comparison of the Danuglipron-bound state versus the Chu-128 state at individual time points (10, 50, 1250, and 6250 seconds). Red signifies peptides with increased HDX between states, while blue represents peptides with decreased HDX. The 98% confidence intervals are shown as black dashed lines, and grey data denote peptides with insignificant ΔHDX. All measurements were performed in triplicate.

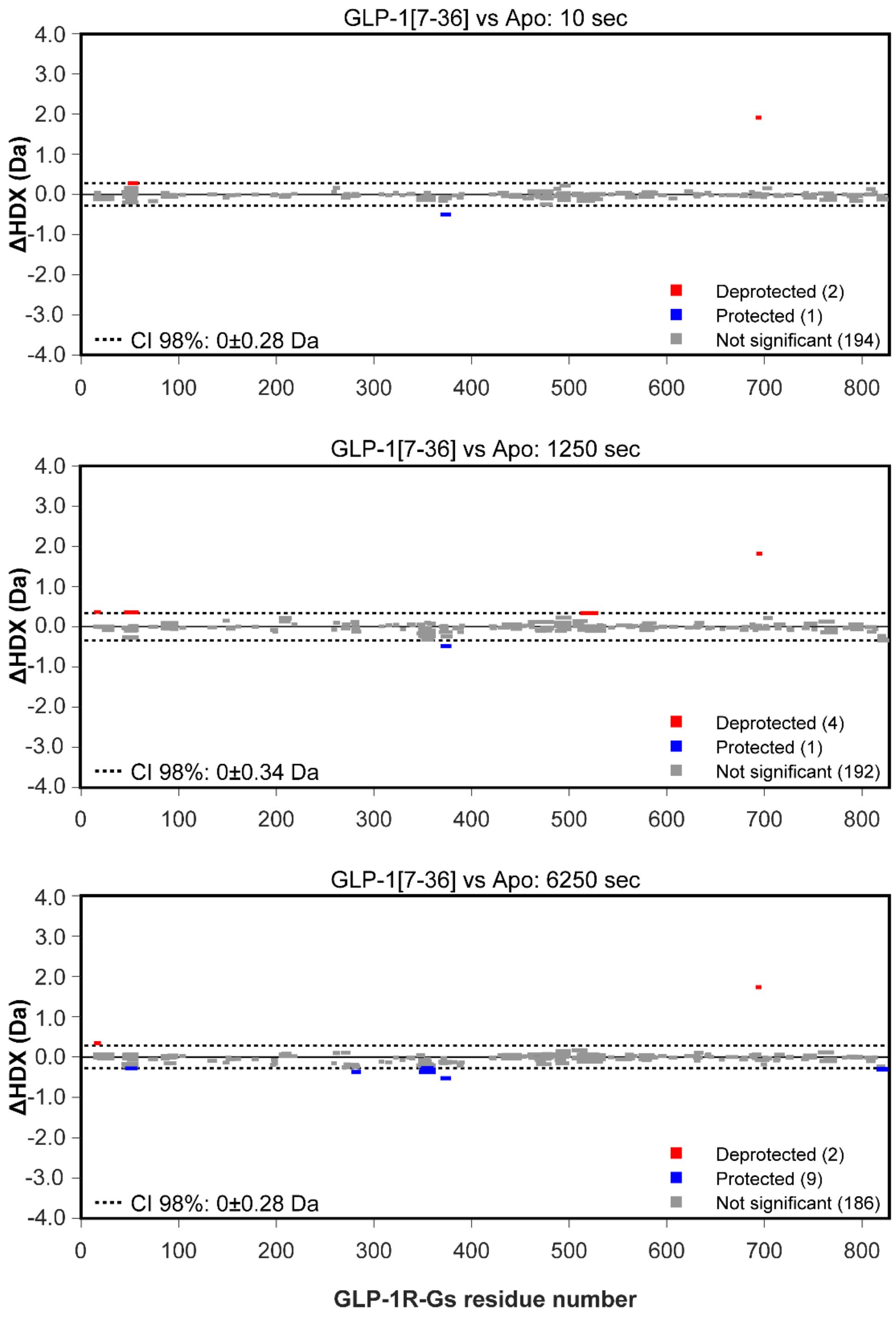


**Figure S9. Individual Woods plots for two-state comparison of GLP-1[7-36] vs Apo.** Differential HDX (ΔHDX) plots from a two-state comparison of the GLP-1[7–36]-bound state versus the apo state at individual time points (10, 1250, and 6250 seconds). Red signifies peptides with increased HDX between states, while blue represents peptides with decreased HDX. The 98% confidence intervals are shown as black dashed lines, and grey data denote peptides with insignificant ΔHDX. All measurements were performed in triplicate.

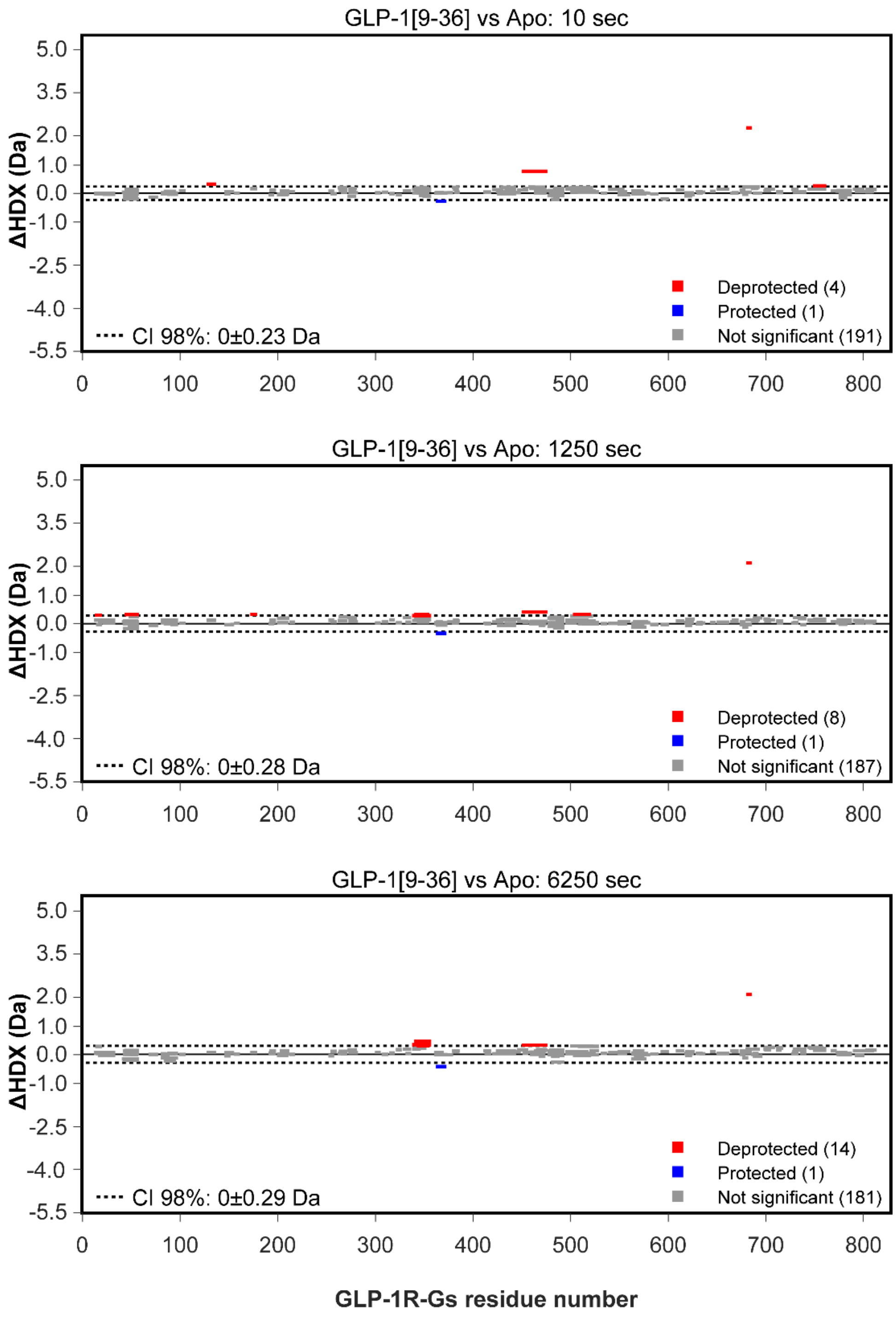


**Figure S10 Individual Woods plots for two-state comparison of GLP-1[9-36] vs Apo.** Differential HDX (ΔHDX) plots from a two-state comparison of the GLP-1[9–36]-bound state versus the apo state at individual time points (10, 1250, and 6250 seconds). Red signifies peptides with increased HDX between states, while blue represents peptides with decreased HDX. The 98% confidence intervals are shown as black dashed lines, and grey data denote peptides with insignificant ΔHDX. All measurements were performed in triplicate.

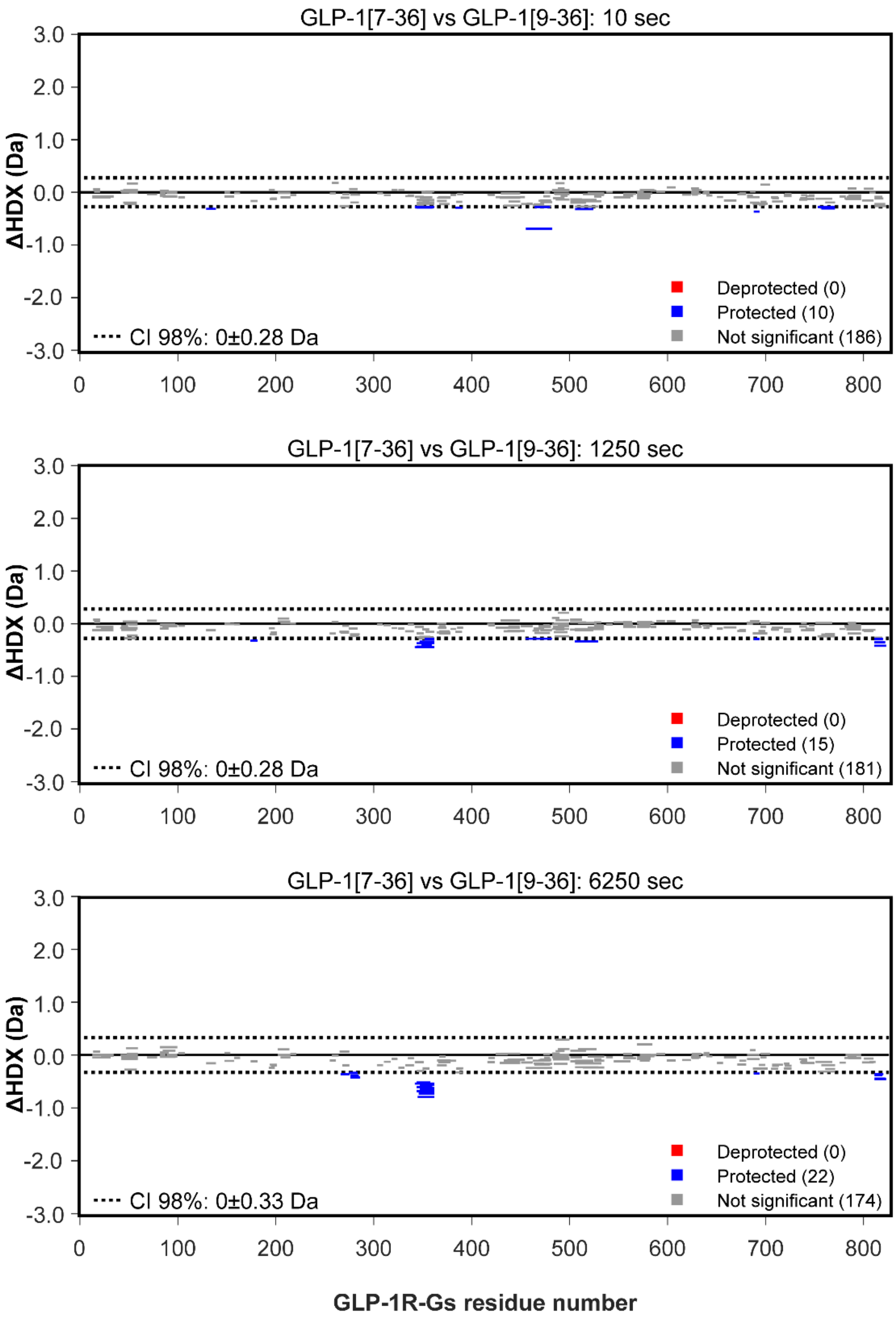


**Figure S11. Individual Woods plots for two-state comparison of GLP-1[7-36] vs GLP-1[9-36].** Differential HDX (ΔHDX) plots from a two-state comparison of the GLP-1[7-36]-bound state versus the GLP-1[9–36]-bound state at individual time points (10, 1250, and 6250 seconds). Red signifies peptides with increased HDX between states, while blue represents peptides with decreased HDX. The 98% confidence intervals are shown as black dashed lines, and grey data denote peptides with insignificant ΔHDX. All measurements were performed in triplicate.

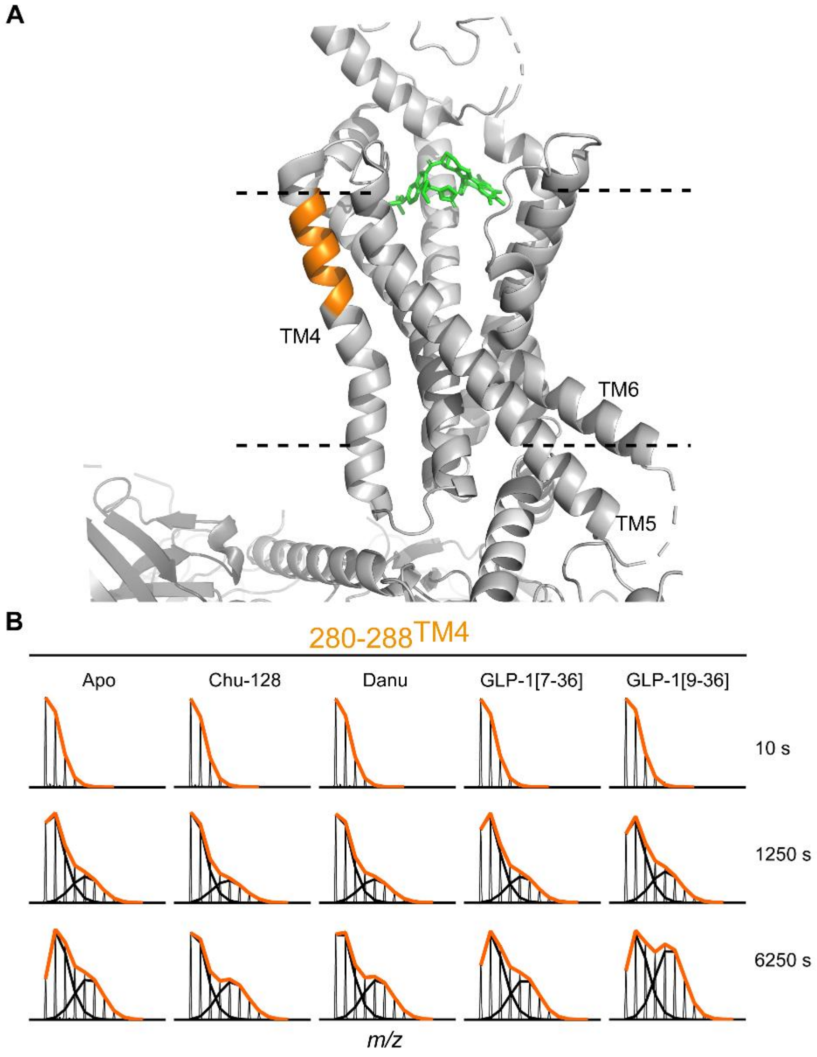


**Figure S12. Allosteric modulation of the TM4 observed in region spanning residues 280-288. (A)** 6XOX pdb structure of GLP-1R in complex with Gs and Orforglipron, with TM4 region of 280-288 highlighted in orange. Complex is positioned relative to the membrane, depicted by two dashed lines, with the Orforglipron moiety shown in green, facing TM4, TM5 and TM6. **(B)** HDX-MS plots showing bimodal distributions in GLP-1R TM4 (residues 280-288) indicative of conformational heterogeneity. Spectra were analysed using HX-Express (v3) and further edited in Prism 9 (version 9.5.1.733). Distinct populations are shown as black lines, and mixed envelopes as lines coloured in orange. The heterogeneity observed in the TM4 region in the apo state is modulated upon GLP-1[9–36] binding, whereas the agonists Chu-128, Danuglipron, and GLP-1[7–36] show no evidence of such modulation.

**Table S1: Human [$^{125}$I]GLP-1(7-36)$NH_2$ binding to human GLP-1R receptor constructs.**

| GLP-1R preparation | FLAG-GLP-1R-Gs fusion + βγ purified protein | FLAG-GLP-1R-Gs fusion + βγ expressed in Sf9 |
|---|---|---|
| GLP-1R protein or membrane capture | Mouse Anti-FLAG IgG + Anti-Mouse IgG PVT-SPA + 0.01% LMNG, 0.001% CHS | WGA-PVT-SPA |
| $B_{max}$ [a] | 0.00621 ± 0.00110 fmol [$^{125}$I]GLP-1(7-36)$NH_2$ bound / fmol of GLP-1R protein | 2320 ± 310 fmol [$^{125}$I]GLP-1(7-36)$NH_2$ bound / mg of Sf9 membrane protein expressing GLP-1R |
| Compound/Peptide | $K_i$ [b] (nM) (SEM, n) | $K_i$ [b] (nM) (SEM, n) |
| GLP-1(7-36)$NH_2$, human | 0.605 (0.092, 8) | 0.279 (0.043, 5) |
| [$^{127}$I]Tyr$^{19}$-GLP-1(7-36)$NH_2$, human | 0.783 [c] (0.068, 8) | 0.195 [c] (0.048, 5) |
| GLP-1(9-36)$NH_2$, human | 174 (32, 8) | 85.6 (14.9, 5) |
| LSN3955529 CHU-128 | 2.34 (0.38, 6) | 0.328 (0.056, 5) |
| LSN3535219 Danuglipron | 135 (22, 6) | 26.0 (3.8, 5) |

[a]$B_{max}$ values were determined by homologous competition and are reported as the arithmetic mean ± SEM.

[b]$K_i$ values reported as the geometric mean with the SEM and the number of independent experiments in parentheses.

[c]$K_d$ values were determined by homologous competition and are reported as the geometric mean with the SEM and the number of independent experiments in parentheses.

**Protein sequences in the apo-active GLP-1R–Gαs–Gβγ complex**

##### > Fusion of GLP-1R (N-CD8 signal peptide–FLAG tag–24–452)–Gαs (26–394, N-term replaced with Gαi1 residues 2–18: GCTLSAEDKAAVERSKM)

MALPVTALLLPLALLLHAARPAAASGIDYKDDDDKRPQGATVSLWETVQKWREYRRQCQRSLTEDPPPATDLFCN
RTFDEYACWPDGEPGSFVNVSCPWYLPWASSVPQGHVYRFCTAEGLWLQKDNSSLPWRDLSECEESKRGERSSPE
EQLLFLYIIYTVGYALSFSALVIASAILLGFRHLHCTRNYIHLNLFASFILRALSVFIKDAALKWMYSTAAQQHQ
WDGLLSYQDSLSCRLVFLLMQYCVAANYYWLLVEGVYLYTLLAFSVLSEQWIFRLYVSIGWGVPLLFVVPWGIVK
YLYEDEGCWTRNSNMNYWLIIRLPILFAIGVNFLIFVRVICIVVSKLKANLMCKTDIKCRLAKSTLTLIPLLGTH
EVIFAFVMDEHARGTLRFIKLFTELSFTSFQGLMVAILYCFVNNEVQLEFRKSWERWRLEHLHIQRDSSMKPLKC
PTSSLSSGATAGSSGCTLSAEDKAAVERSKMIEKQLQKDKQVYRATHRLLLLGAGESGKNTIVKQMRILHVNGFN
GEGGEEDPQAARSNSDGEKATKVQDIKNNLKEAIETIVAAMSNLVPPVELANPENQFRVDYILSVMNVPDFDFPP
EFYEHAKALWEDEGVRACYERSNEYQLIDCAQYFLDKIDVIKQADYVPSDQDLLRCRVLTSGIFETKFQVDKVNF
HMFDVGAQRDERRKWIQCFNDVTAIIFVVASSSYNMVIREDNQTNRLQAALKLFDSIWNNKWLRDTSVILFLNKQ
DLLAEKVLAGKSKIEDYFPEFARYTTPEDATPEPGEDPRVTRAKYFIRDEFLRISTASGDGRHYCYPHFTCSVDT
ENIRRVFNDCRDIIQRMHLRQYELL*

##### > MAL-Gβ1(2–340)

MALSELDQLRQEAEQLKNQIRDARKACADATLSQITNNIDPVGRIQMRTRRTLRGHLAKIYAMHWGTDSRLLVSA
SQDGKLIIWDSYTTNKVHAIPLRSSWVMTCAYAPSGNYVACGGLDNICSIYNLKTREGNVRVSRELAGHTGYLSC
CRFLDDNQIVTSSGDTTCALWDIETGQQTTTFTGHTGDVMSLSLAPDTRLFVSGACDASAKLWDVREGMCRQTFT
GHESDINAICFFPNGNAFATGSDDATCRLFDLRADQELMTYSHDNIICGITSVSFSKSGRLLLAGYDDFNCNVWD
ALKADRAGVLAGHDNRVSCLGVTDDGMAVATGSWDSFLKIWN*

##### > 8×His-Gγ2(2–70)

MHHHHHHHHLVPRGSASNNTASIAQARKLVEQLKMEANIDRIKVSKAAADLMAYCEAHAKEDPLLTPVPASENPF
REKKFFCAIL*
